# Remodeling of the Ribosomal Quality Control and Protein Translation by a Viral Ubiquitin Deconjugase

**DOI:** 10.1101/2023.01.31.526464

**Authors:** Jiangnan Liu, Noemi Nagy, Francisco Aguilar-Alonso, Francisco Esteves, Carlos Ayala-Torres, Shanshan Xu, Maria G. Masucci

## Abstract

The strategies adopted by viruses to reprogram the protein translation and quality control machineries to promote infection are poorly understood. Here, we discovered that the viral ubiquitin deconjugase (vDUB) encoded in the large tegument protein of Epstein-Barr virus (EBV) regulates ribosomal stress responses. The vDUB participates in protein complexes that include the ubiquitin ligases ZNF598 and LTN1 and the UFM1 ligase UFL1. Upon ribosomal stalling, the vDUB counteracts the ubiquitination of 40S ribosome subunits, inhibits the degradation of translation-stalled polypeptides by the proteasome, and prevents UFMylation of the 60S particle, which impairs the ER-phagy- dependent clearance of stalled products. Inhibition of the ribosome quality control activates a GCN2-dependent integrated stress response that decreases global protein translation while promoting the readthrough of stall-inducing mRNAs. The vDUB enhances viral mRNAs translation and virus release during productive infection, pointing to a pivotal role in cell reprogramming that enables virus production and underlies the pathogenesis of EBV-associated cancers and autoimmune diseases.

**GRAPHIC SUMMARY:** 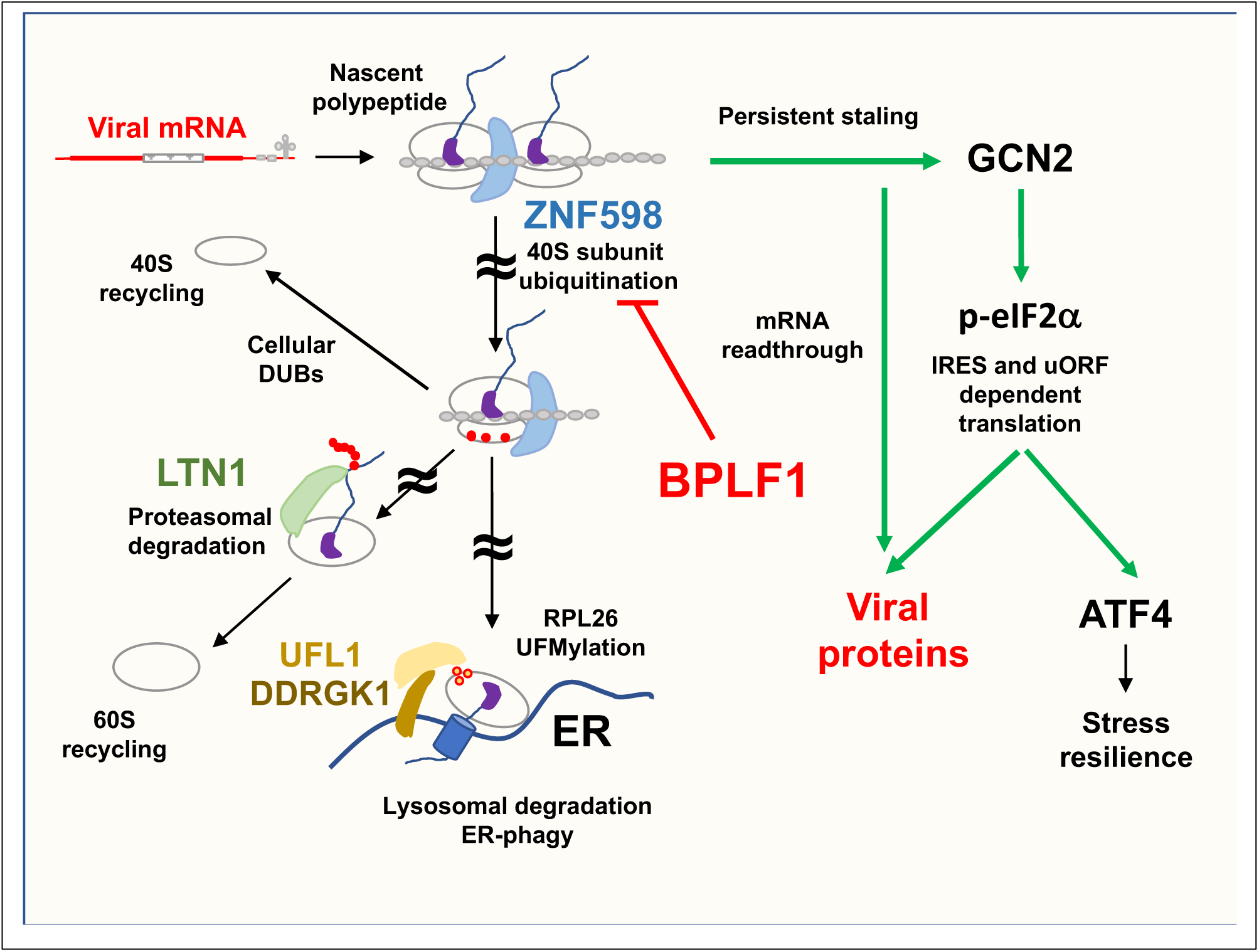

## INTRODUCTION

Translating viral mRNAs is particularly challenging due to the presence of long repetitive sequences, complex secondary structures, suboptimal codon usage, and the frequent occurrence of nucleotide misincorporations that may slow translation and induce ribosome stalling (Chyzynska et al., 2021). While pausing the elongation cycle may resolve some issues, persistent stalling causes ribosome collision that may disrupt protein synthesis and ultimately cause cell death (Vind et al., 2020). To rescue stalled ribosomes, the cell deploys a ribosome-associated quality control (RQC) machinery that recognizes collided di- and trisomes (Ikeuchi et al., 2019) and triggers site-specific mono- ubiquitination of the 40S particle (Garzia et al., 2021; Jung et al., 2017; Juszkiewicz and Hegde, 2017; Kim et al., 2018; Matsuo et al., 2017). In mammals, ubiquitination of the RPS10 (eS10) and RPS20 (uS10) subunits by ZNF598 (Garzia et al., 2017; Juszkiewicz et al., 2018; Juszkiewicz and Hegde, 2017; Matsuo et al., 2017; Sundaramoorthy et al., 2017) promotes ribosome disassembly (Juszkiewicz et al., 2020; Matsuo et al., 2020), followed by the extraction of nascent polypeptides, their ubiquitination by Listerin-1 (LTN1), and degradation by the proteasome (Bengtson and Joazeiro, 2010; Brandman et al., 2012). This allows the recycling of ribosome particles, while ubiquitination of RPS2 (uS5) and RPS3 (uS3) by ZNF598 and RNF10 (Garzia et al., 2021) targets irreversibly stalled ribosomes for lysosomal degradation (Garzia et al., 2021; Meyer et al., 2020). Although potentially deadly, failure of the RQC may also promote translation since a prolonged pause may facilitate the readthrough of stall-inducing mRNAs (Sundaramoorthy et al., 2017).

A corresponding pathway featuring post-translational modification of the 60S particle by the ubiquitin-like modifier-1 (UFM1) rescues ribosomes that stall while synthesizing proteins destined for the endoplasmic reticulum (ER) (Gerakis et al., 2019; Witting and Mulder, 2021). UFMylation of RPL26 (uL24) by the ER-associated UFL1/DDRGK1/CDK5RAP3 ligase complex (Liang et al., 2020; Peter et al., 2022; Stephani et al., 2021; Tatsumi et al., 2010) initiates a signaling cascade leading to the lysosomal degradation of translocon-trapped polypeptide together with a portion of the ER in a process known as ER-phagy (Ferro-Novick et al., 2021; Walczak et al., 2019; Wilkinson, 2019). Failure of this pathway upon functional inactivation of the ligase (Cai et al., 2015; Tatsumi et al., 2010; Walczak et al., 2019) triggers the unfolded protein response and reprograms protein synthesis in favor of cap-independent, internal ribosomal entry site (IRES)- or upstream open reading frame (uORF)-dependent translation (Komar and Hatzoglou, 2011; Ruggiano et al., 2014; Starck et al., 2016). The challenging structure of viral mRNA and the depletion of tRNAs induced by the mass production of comparatively small proteomes (Nunes et al., 2020) suggest that viruses may be exquisitely dependent on the RQC and ER-RQC for efficient protein production.

Epstein-Barr virus (EBV) is a lymphotropic herpesvirus that establishes persistent infections in most human adults and participates in the pathogenesis of lymphoid and epithelial cell malignancies (Shannon-Lowe and Rickinson, 2019). Increased levels of circulating virus and specific antibodies precede the clinical manifestation of EBV-associated malignancies and non-malignant autoimmune conditions (Houen and Trier, 2020; Inagaki et al., 2020; Munz, 2021) pointing to an important role of the productive virus cycle in disease pathogenesis. During productive EBV infection, more than seventy different viral proteins reprogram the cellular environment to allow virus replication and provide the building blocks of new virus particles (Kenney, 2007). The known viral mechanisms for interfering with translation often involve the expression of RNA-binding proteins that mimic or hijack components of the translation machinery (Daly et al., 2020). Here, we have discovered a new strategy by which EBV reprograms translation by harnessing the activity of the ubiquitin deconjugase encoded in the N-terminal domain of the large tegument protein BPLF1 (vDUB) to inhibit the RQC and ER-RQC, trigger a GCN2 kinase-dependent integrated stress response, and promote the readthrough of stall-inducing viral mRNAs.

## RESULTS

### BPLF1 participates in protein complexes involved in translation and ribosomal quality control

The Gene ontology (GO) classification of proteins that co-immunoprecipitate with the vDUB produced by caspase-1 cleavage of the N-terminus (aa 1-235) of the EBV large tegument protein BPLF1 (Gastaldello et al., 2013) identified a broad range of cellular functions underlying a pleiotropic role in host-cell remodeling (Gupta et al., 2018). Analysis of the enriched GO classes revealed the enrichment of proteins involved in regulating RNA metabolism, ribosome biogenesis, mRNA translation, and proteotoxic cell responses (Figure 1A). STRING analysis of the protein interaction network identified a major hub centered around the ribosome and components of the mRNA translation, ER trafficking, and ER stress response machineries (Figure 1B and S1 Table). The enriched proteins included the translation pre-initiation complex subunit eIF2α that, upon phosphorylation, becomes an inhibitor of global protein synthesis (Gebauer and Hentze, 2004), the translation initiation complex subunits eIF4G1 (Gingras et al., 1999) and eIF3E (Masutani et al., 2007), and the ribosome pause release factor eIF5A (Pelechano and Alepuz, 2017); several subunits of the 40S and 60S ribosome particles; the SRP68 subunit of the Signal Recognition Particle (SRP) and the β subunit of the SRP receptor (SRPRB) (Akopian et al., 2013); the SEC63 subunit of the ER translocon (Osborne et al., 2005), and the Ribophorin (RPN)-1 subunit of the translocon-associated N-oligosaccharyltransferase (OST) complex (Kelleher and Gilmore, 2006). The interaction network also included the RQC ligase ZNF598, the AAA+ ATPase VCP/p97 that extracts ubiquitinated nascent polypeptides from stalled ribosomes (Verma et al., 2013), and the UFM1 ligase UFL1 that regulates the ER-RQC and ER-phagy responses (Banerjee et al., 2020; Witting and Mulder, 2021) (Figure 1B). Several members of the interaction network, including RPL26 (Walczak et al., 2019), RPN1 (Liang et al., 2020), SRPRB, eIF5A, eIF2α and eIF4G1 (Gak et al., 2020) (highlighted by red circles in Figure 1B), are putative UFL1 substrates suggesting that BPLF1 may interfere with UFMylation-regulated processes.

**Figure 1.**
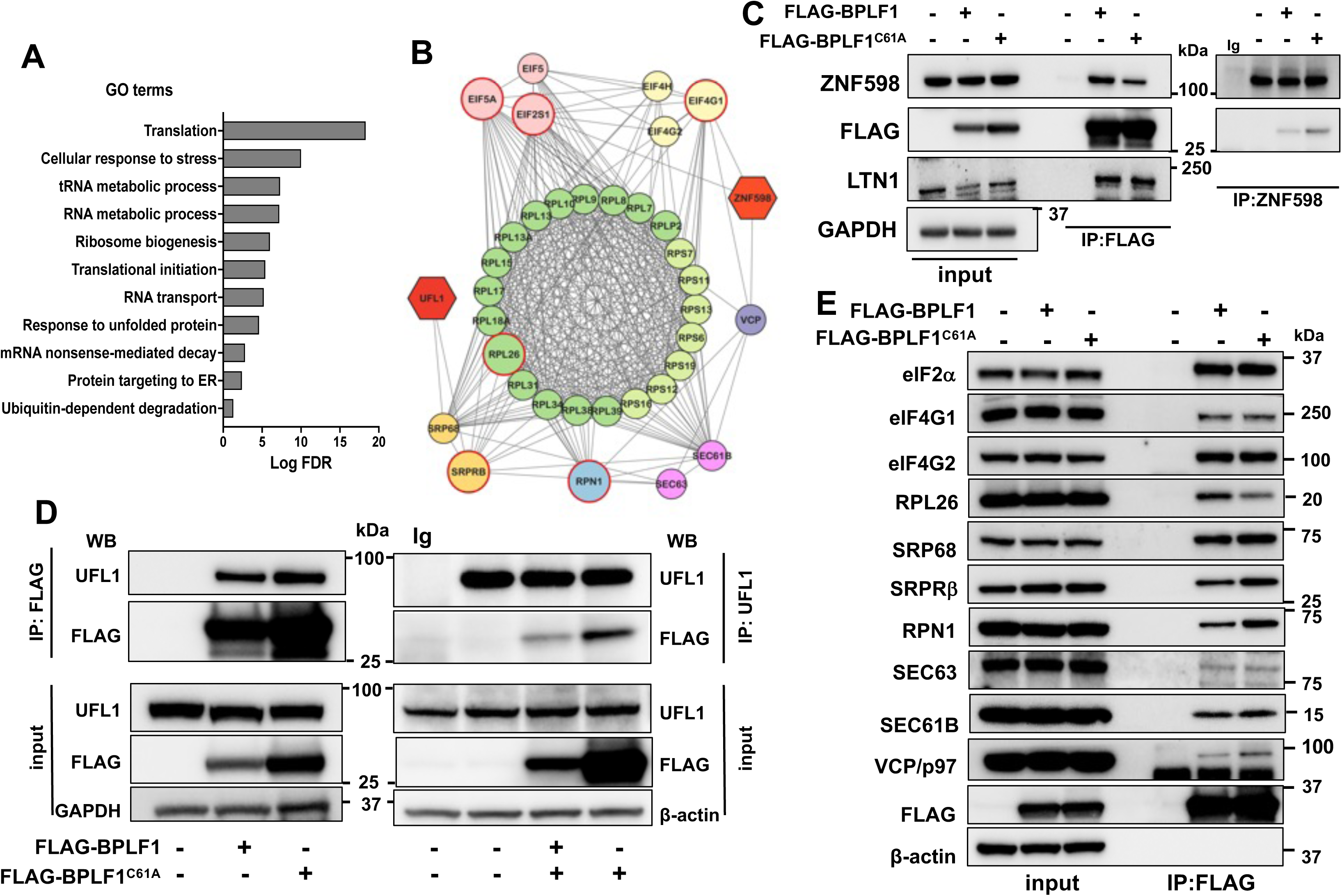
BPLF1 interacts with proteins involved in translation and ribosomal stress responses. (**A**) Significantly enriched biological processes in the BPLF1 interactome. One hundred sixty-nine out of 375 proteins exhibiting fold enrichment ≥2 (45% of the interactome) were annotated in these processes. (**B**) STRING network diagram of the major protein interaction hub including several ribosome subunits, components of the mRNA translation, ER trafficking, and ER stress responses. The functional annotation is color-coded: light green, 40S ribosome subunit; dark green 60S ribosome subunit; pink, translation pre-initiation complex; light yellow, translation initiation complex; dark yellow, signal recognition particle (SRP) and SRP receptor (SRPB) involved in the recognition and translocation of signal-sequence-tagged proteins; magenta, ER translocon complex that mediates forward and retrograde transport across the ER; light blue, translocon-associated N-oligosaccharyltransferase (OST) complex that links high mannose sugars to the Asn-X-Ser/Thr consensus motif of ER-translocated polypeptides; violet, AAA+ ATPase VCP/p97 that participated in the extraction of ubiquitinated polypeptides from protein complexes; red, ubiquitin and UFM1 ligases. Red borders highlight previously described putative UFMylation substrates. (**C**) Western blots illustrating the interaction of BPLF1 with the ZNF598 and LTN1 ubiquitin ligases. Cell lysates of HEK293T cells transfected with plasmids expressing FLAG- BPLF1/BPLF1^C61A^ or the empty FLAG vector were immunoprecipitated with either anti-FLAG coated beads or antibodies to ZNF598 followed by capture with ProtA-coated beads. An isotype-matched Ig control was included in the ZNF598 immunoprecipitation to verify specificity. Western blots were probed with the indicated antibodies. Blots from one representative experiment out of two are shown in the figure. (**D**) Western blots illustrating the interaction of BPLF1 with the UFM1 ligase UFL1. Blots from one out of two independent experiments are shown in the figure. (**E**) Representative western bots illustrating the interaction of BPLF1 with a selection of the proteins identified in B. Each interaction was validated in at least two independent co-immunoprecipitation experiments.

The interactions were validated by co-immunoprecipitation in cell lysates of HEK293T cells transfected with FLAG-BPLF1, the catalytic mutant FLAG-BPLF1^C61A^, or FLAG-empty vector. ZNF598 and UFL1 were readily detected in the BPLF1 immunoprecipitates (Figure1C, 1D). Conversely, BPLF1 was found in the ZNF598 and UFL1 immunoprecipitates, confirming that the vDUB interacts with the ligases of the RQC and ER-RQC pathways. Probing the FLAG immunoprecipitates with a selection of antibodies specific to other members of the protein interaction network, confirmed the presence of BPLF1 in protein complexes involved in translation and the ER translocation and glycosylation of nascent polypeptides (Figure 1E). Although they did not meet the mass spectrometry significance threshold, LTN1 (Figure 1D) and the SEC61B component of the ER translocon (Figure 1E) were regularly detected in the immunoprecipitates.

### BPLF1 inhibits the activation of RQC and stabilizes RQC substrates

Upon sensing collided ribosomes, ZNF598 initiates the RQC by ubiquitinating RPS10 and RPS20, while ubiquitination of RSP2 and RPS3 by ZNF598 and RNF10 promotes lysosomal degradation of the 40S particle (Garzia et al., 2021). To probe the effect of the vDUB on these processes, we first tested whether it affects the ubiquitination of 40S subunits in cells treated with the translation elongation inhibitor Anisomycin (ANS). Control and ANS-treated FLAG-BPLF1/BPLF1^C61A^ transfected HEK293T cells were lysed in a buffer containing NEM and iodoacetamide to inhibit DUB activity. Band shifts corresponding to the size of mono-ubiquitinated RPS10 and RPS3 were readily detected in the western blots of ANS-treated cells (Figure 2A). The expression of BPLF1 significantly impaired the appearance of ubiquitinated species. In contrast, the catalytic mutant BPLF1^C61A^ had no consistent effect (Figure 2B). To test whether subsequent steps of the RQC pathway are affected, we took advantage of a GFP-reporter lacking the termination codon, GFP-nonSTOP (Figure 2C). The progression of translation through the poly(A) tail leads to ribosome stalling, activation of the RQC, ubiquitination of the C-terminally extended product by LTN1 and degradation by the proteasomes (Akimitsu et al., 2007; Filbeck et al., 2022). Western blots of HEK293T cells transfected with the GFP-nonSTOP reporter alone or with FLAG-BPLF1/BPLF1^C61A^ were probed with antibodies to GFP. As a control for proteasomal degradation, the cells were treated with MG132 during the last 6 h before harvesting. A weak GFP band was detected in cells transfected with the GFP-nonSTOP plasmid alone, while treatment with MG132 promoted the accumulation of a slightly larger C-terminally extended product (Figure 2D). An even more substantial accumulation of the extended product was observed in BPLF1 expressing cells while the BPLF1^C61A^ mutant had no effect (Figure 2E), supporting the conclusion that the active enzyme inhibits the RQC and prevents the degradation of translation-arrested polypeptides.

**Figure 2.**
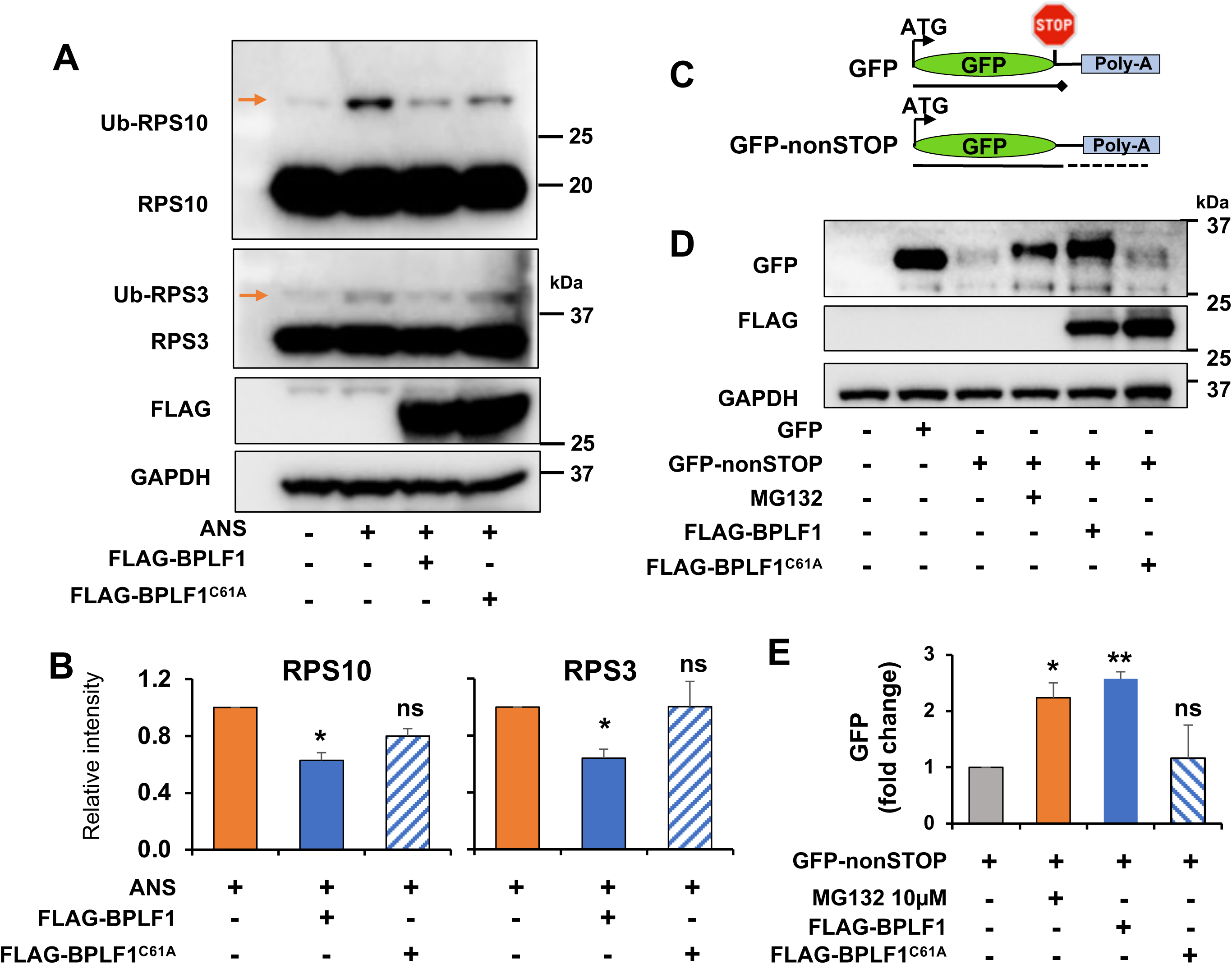
BPLF1 inhibits the activation of RQC in ANS-treated cells and rescues RQC substrates from proteasomal degradation. (**A**) BPLF1 inhibits the ubiquitination of 40S ribosome subunits in ANS treated cells. HeLa cells transfected overnight with plasmids expressing FLAG-tagged BPLF1, the BPLF1^C61A^ mutant, or the empty FLAG vector were treated with 0.5 μg/ml ANS for 30 min and then lysed in buffer containing NEM and iodoacetamide to inhibit DUB activity. Western blots were probed with the indicated antibodies. Blots from one representative experiment out of three are shown in the figure. (**B**) The intensity of the Ub-RPS10 and Ub-RPS3 bands was quantified by densitometry in three independent experiments. The relative intensity of the bands in transfected versus control cells is shown. Significance was calculated by Student t-test. *P= ≤0.05. (**C**) Schematic illustration of the GFP-nonSTOP reporter. Following deletion of the stop codon, translation through the poly(A) tail leads to ribosome stalling, activation of the RQC, and degradation of the C-terminally extended product by the proteasomes (**D**). Western blots of HEK293T cells transfected with a control GFP plasmid or the GFP-nonSTOP reporter alone or with FLAG-BPLF1/BPLF1^C61A^ were probed with antibodies to GFP. As a control for proteasomal degradation, the GFP-non-Stop transfected cells were treated with 10 μM MG132 during the last 6 h before harvesting. Western blots from one representative experiment out of two are shown in the figure. (**E**). The intensity of the GFP band was quantified by densitometry in two independent experiments. The relative intensity of the bands in GFP-nonSTOP in MG132-treated and transfected cells versus control cells is shown. Significance was calculated by Student t-test. * = P≤0.05, ** = P≤0.01

### BPLF1 inhibits RPL26 UFMylation by interfering with activation of the UFL1 ligase

Ribosome stalling at the ER triggers the UFMylation of RPL26, which promotes the lysosomal degradation of translocon-trapped polypeptides and the activation of ER-phagy (Ferro-Novick et al., 2021; Liang et al., 2020; Walczak et al., 2019; Wang et al., 2020). To investigate whether the vDUB interferes with RPL26 UFMylation, HeLa cells transfected with Myc-tagged RPL26 alone or with FLAG-BPLF1/BPLF1^C61A^ were treated with ANS followed by lysis under denaturing conditions to destroy non-covalent interactions, immunoprecipitation with anti-Myc coupled beads and western blot analysis. Three bands of size corresponding to mono-, di- and tri-UFMylated RPL26 were detected by probing the blots with antibodies to UFM1 (Figure 3A). Two bands correspond to previously described UFMylation sites on Lys134 and Lys132 (Walczak et al., 2019), while the third band may indicate poly-UFMylation. The UFMylated species were virtually absent in cells expressing catalytically active BPLF1, with corresponding accumulation of unmodified RPL26, while the catalytic mutant had no consistent effect. The UFMylation of endogenous RPL26 was significantly impaired in FLAG-BPLF1 transfected HeLa cells treated with ANS (Figure 3B and 3C). Of note, a band of size corresponding to mono-UFMylated RPL26 was regularly detected by the UFM1-specific antibody in the untreated cells, independently of BPLF1 expression. This finding is in line with the observations that UFMylated RPL26 is incorporated in translating ribosomes at steady state and that UFMylation of RPL26-K134 is a prerequisite for the UFMylation of additional sites (Walczak et al., 2019). Comparable inhibition of RPL26 UFMylation was observed when HeLa cells were co-transfected with BPLF1 and an ER-targeted ribosomal stalling reporter (Figure S1A, S1B), confirming the capacity of the vDUB to halt the UFMylation of the endogenous protein upon induction of ribosomal stress.

**Figure 3.**
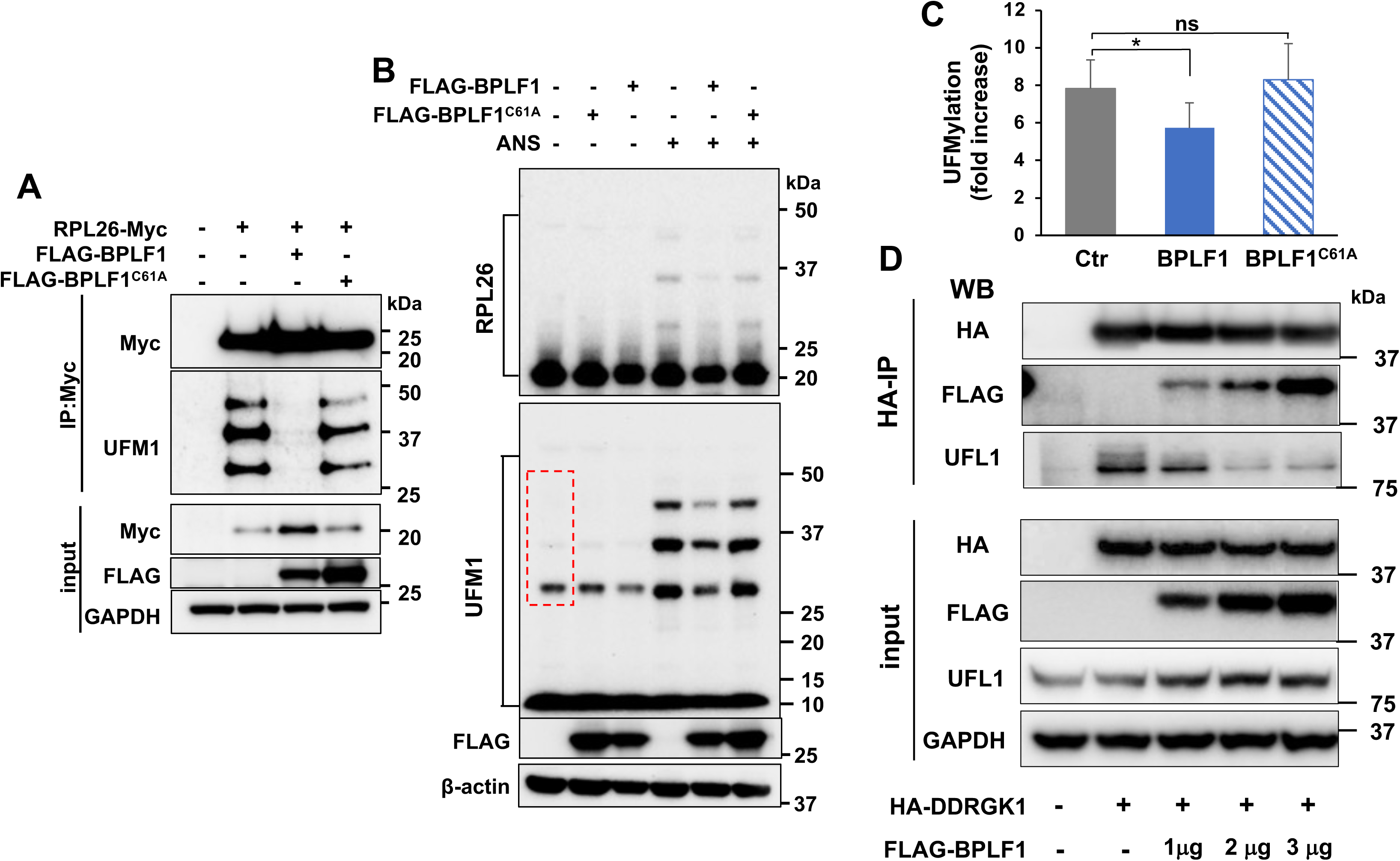
BPLF1 inhibits UFMylation and the binding of UFL1 to DDRGK1. (**A**). BPLF1 halts the UFMylation of RPL26 in ANS-treated cells. HeLa cells transfected with Myc-tagged RPL26 alone or with FLAG-BPLF1/BPLF1^C61A^ were treated with 50 ng/ml ANS for 1 h, followed by lysis under denaturing conditions to destroy non-covalent interactions and immunoprecipitation with anti-Myc coupled beads. Western blots from one representative experiment out of four are shown in the figure. (**B**) BPLF1 inhibits the UFMylation of endogenous RPL26 in ANS-treated cells. Control and FLAG- BPLF1/BPLF1^C61A^ transfected HeLa cells were cultured for 24 h and then treated with 50 ng/ml ANS for 1 h before lysis in buffer containing NEM and iodoacetamide to inhibit DUB activity. High molecular weight species corresponding to the size of mono- di- and tri-UFMylated RPL26 were detected in western blots probed with antibodies specific for RPL26 and UFM1. Representative blots from one out of three independent experiments are shown in the figure. (**C**) Densitometry quantification of the intensity of the UFMylated species indicated by a red dotted box. The mean ± SD fold increase in ANS treated relative to untreated cells in three independent experiments is shown. (**D)** BPLF1 inhibits the interaction of the UFL1 ligase with the ER-anchoring subunit DDRGK1. HeLa cells were co-transfected with plasmids expressing HA-DDRGK1 and increasing amounts of Flag-BPLF1, followed by immunoprecipitation with anti-HA-tag beads. Western bots were probed with the indicated antibodies. Representative blots from one out of three independent experiments are shown in the figure.

To test whether, in addition to the known specificity for ubiquitin and Nedd8 conjugates (Gastaldello et al., 2010), the viral enzyme may also have de-UFMylase activity, we performed *in vitro* assays where recombinant BPLF1 and the cellular deUFMylase UFSP2 were tested for their effect on the UFMylation of Histone H3/H4 by purified components of the UFMylation cascade (Figure S2). UFMylated H3 was readily detected when the reaction was performed in the presence of ATP. Recombinant BPLF1 had no appreciable effect, whereas small amounts of UFSP2 substantially decreased UFMylation. Similar results were obtained when increasing amounts of recombinant BPLF1 were tested for the capacity to hydrolyze UFM1-GFP and Ub-GFP reporters. While the Ub-GFP reporter was efficiently cleaved by small quantities of BPLF1, the UFM1-GFP reporter was hardly affected even by the highest amount of BPLF1 tested (Figure S3A). To explore the possibility that critical co-factors may be missing in the *in vitro* reaction, a reporter plasmid expressing a UFM1-GFP fusion protein was co-transfected in U2OS cells lacking the endogenous UFSP2 (Figure S3B), together with plasmids expressing FLAG-BPLF1 or FLAG-UFSP2 (Figure S3C). The reporter was efficiently cleaved by UFSP2, resulting in the accumulation of free GFP and UFM1, whereas BPLF1 had no effect.

The lack of deUFMylase activity suggests that the UFL1 ligase may be the primary target of the inhibitory effect. UFL1 is recruited to the ER by binding to DDRGK1, which is also required for optimal ligase activity (Peter et al., 2022). Therefore, we asked whether BPLF1 interferes with this interaction. A plasmid expressing HA-tagged DDRGK1 was co-transfected in HeLa cells with increasing amounts of the FLAG-BPLF1 plasmid, and the formation of DDRGK1-UFL1 complexes was assayed by probing Myc immunoprecipitates with a UFL1 specific antibody. BPLF1 inhibited the recruitment of endogenous UFL1 to DDRGK1 in a dose-dependent manner (Figure3D), pointing to the failure to assemble the active ligase as the primary inhibitory event. Notably, the inability of catalytic mutant BPLF1^C61A^ to inhibit UFMylation (Figure 3A, 3B) despite efficient binding to UFL1 (Figure 1C) suggests that a ubiquitination-driven triggering event may be required for the assembly of the ligase in cells.

### BPLF1 stabilizes ER-RQC substrates and inhibits ER-phagy

We then investigated whether BPLF1 may regulate the downstream effects of RPL26 UFMylation. To test whether BPLF1 interferes with the degradation of translocon- trapped polypeptides, we took advantage of a ribosome stalling reporter that expresses in-frame an N-terminal signal sequence followed by an N-glycosylation site, GFP, and a stretch of 20 Lys residues encoded by AAA codons (ER-K20) (Figure 4A). Weak bands of approximately 46 kDa and 48 kDa, corresponding to the expected size of glycosylated ER-K20 polypeptides, and a band of about 50 kDa, were detected in transfected HEK293T cells (Figure 4B). The intensity of the bands was strongly increased in cells expressing BPLF1 or treated with Bafilomycin A1 (BafA1), that prevents the acidification of lysosomes (Bowman et al., 1988). In contrast, expression of the BPLF1^C61A^ mutant or treatment with the proteasome inhibitor MG132 had no significant effect (Figure 4C), supporting the conclusion that vDUB inhibits the lysosomal degradation of translocon-trapped polypeptides.

**Figure 4.**
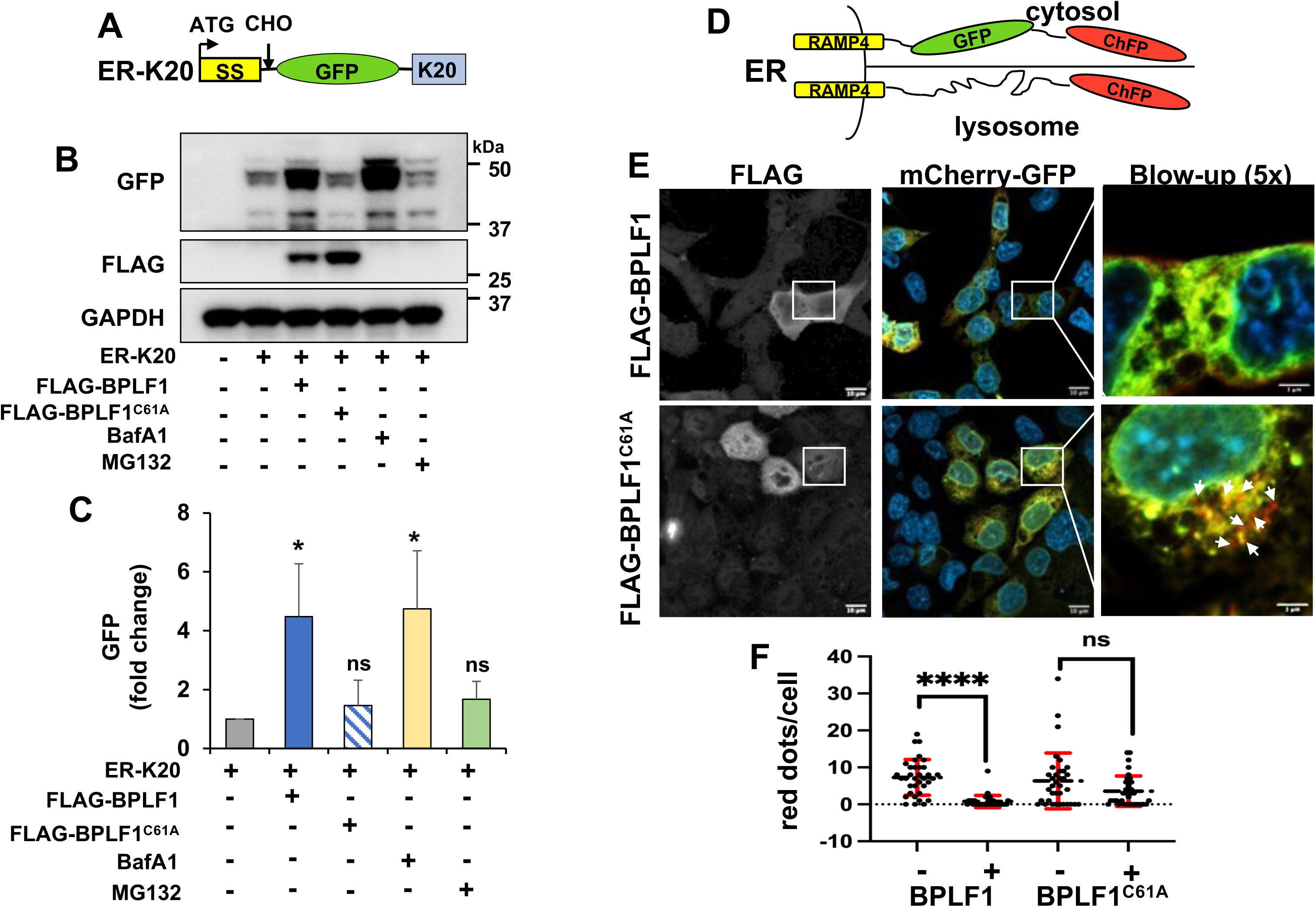
BPLF1 rescues ER-RQC substrates and inhibits ER-phagy. (**A**). Schematic illustration of the ER-RQC reporter. The ER-K20 reporter expresses in-frame an N-terminal ER-targeting signal sequence followed by an N-glycosylation site, GFP, and a stretch of Lys residues encoded by AAA codons (K20). Stalling of the ribosome at the poly(A) sequence traps the nascent ER-inserted polypeptide in the translocon, which promotes lysosome-dependent degradation. (**B**). BPLF1 stabilizes an ER-RQC substrate. HEK293T co-transfected with the ER-K20 reporter and FLAG-BPLF1/BPLF1^C61A^ or the empty FLAG vector were cultured for 24 h. Aliquots of Vector-ER-K20 transfected cells were treated with 0.1 μM BafA1 or 10 μM MG132 during the last 6 h before harvesting. Western blots from one representative experiment out of three are shown in the figure. (**C**). Densitometry quantification of the GFP bands in three independent experiments. Fold increase was calculated as the ratio between the intensity of the band in treated cells versus cells transfected with the ER-K20 reporter alone. Significance was calculated by Student t-test. * = P≤0.05. (**D**). Schematic illustration of the ER-Autophagy Tandem Reporter (EATR). The reporter expresses in-frame the coding sequence of the RAMP4 subunit of the ER translocon complex followed by the coding sequences of eGFP and ChFP. Upon ER insertion of the RAMP4 domain, eGFP and ChFP face the cytosol and emit equal fluorescence, whereas, due to the selective loss of eEGP fluorescence at low pH, ER-loaded autophagosomes appear as distinct red fluorescent dots. (**E**) Representative confocal images illustrating the failure to accumulate red fluorescent dots in cells expressing BPLF1. Stable HCT116-EATR cells were transfected with plasmids expressing FLAG-BPLF1/BPLF1^C61A^ and then starved overnight in EBSS medium before visualizing the formation of ER-loaded autophagosomes by confocal microscopy. (**F**). Quantification of the number of red fluorescent dots in BPLF1/BPLF1^C61A^ positive and negative cells from the same transfection experiments. The cumulative data from two independent experiments where approximately 50 vDUB positive and negative cells were scored are shown. Significance was calculated by Student t-test. **** = P≤0.0001.

The UFMylation of RPL26 and Ribophorin 1 (RPN1) triggers ER-phagy (Liang et al., 2020). To test whether BPLF1 impacts this process, we produced a subline of HCT116 expressing a Dox-regulated ER-phagy reporter constructed by in-frame fusion of the coding sequences of the RAMP4 subunit of the ER translocon, enhanced GFP (eGFP), and Cherry fluorescent protein (ChFP) (Liang et al., 2018) (Figure 4D and Figure S4A). Upon ER insertion of the RAMP4 domain, eGFP and ChFP face the cytosol and emit equal fluorescence, whereas, due to the selective loss of eGFP fluorescence at low pH, ER-loaded autophagosomes appear as distinct red fluorescent dots (Figure S4B). The reporter cell line was transfected with plasmids expressing FLAG-BPLF1/BPLF1^C61A^ in the presence of Dox and then starved overnight in EBSS medium before visualizing the formation of phagolysosomes by confocal microscopy. Red cargo-loaded phagolysosomes were readily detected in cells expressing the BPLF1^C61A^ mutant but not in cells expressing the active BPLF1 (Figure 4E). Quantification of the number of red dots in BPLF1/BPLF1^C61A^ positive and negative cells from the same transfection revealed a highly significant suppression of ER-phagy in cells expressing the catalytically active vDUB (Figure 4F).

### BPLF1 activates a GCN2-dependent Integrated Stress Response (ISR)

Failure of the RQC and ER-RQC activates stress kinases that converge on phosphorylation of the translation initiation factor eIF2α on Ser51 (Donnelly et al., 2013). The ISR promotes cell survival by reducing global protein synthesis while favoring the translation of mRNAs that can bypass the inhibitory effect of eIF2α phosphorylation (Lu et al., 2004). To investigate whether BPLF1 regulates the ISR, we monitored the phosphorylation of eIF2α and the upregulation of ATF4 and its transcriptional target CHOP in cells transfected with FLAG-BPLF1/BPLF1^C61A^ or treated with the ER stress inducer Thapsigargin (TPG) (Sehgal et al., 2017). A comparable upregulation of p-eIF2α, ATF4, and CHOP was observed in TPG-treated and BPLF1-expressing cells, while the BPLF1^C61A^ mutant had no consistent effect (Figure 5A, 5B). In line with the induction of eIF2α phosphorylation, the expression of BPLF1 was accompanied by a highly significant global decrease of mRNA translation as measured by puromycin incorporation assays (Figure S5A, S4B). This was confirmed at the single-cell level by FLAG and puromycin double staining, which revealed a substantial decrease of puromycin staining in cells expressing BPLF1 but not in cells expressing the BPLF1^C61A^ mutant (Figure S5C, S5D).

**Figure 5.**
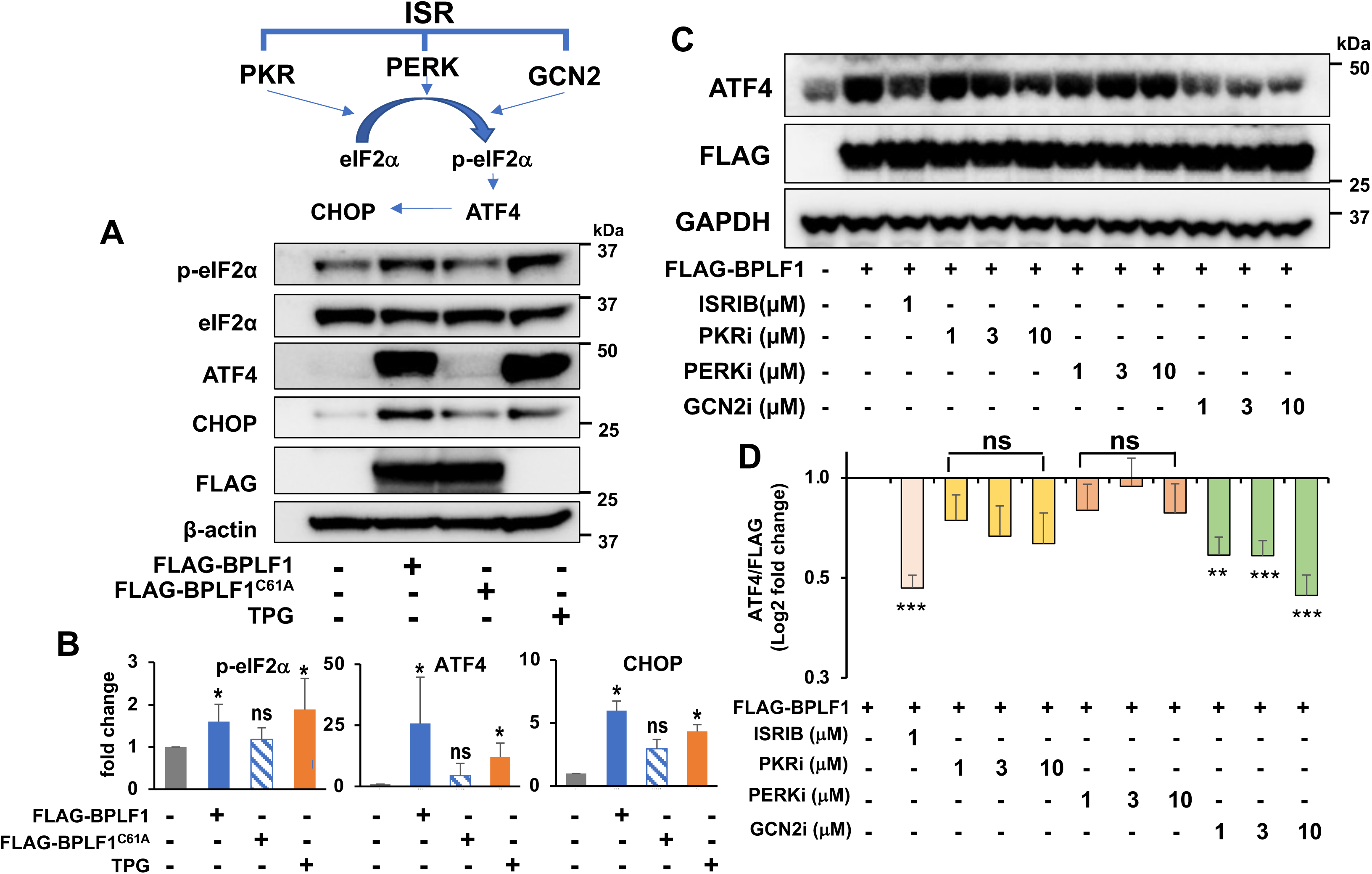
BPLF1 triggers a GCN2-dependent ISR. (**A**). HeLa cells transfected with FLAG-BPLF1/BPLF1^C61A^ or the empty FLAG vector and cultured overnight. The vector-transfected cells were treated with 1 μM of the ER stress inducer Sarco/endoplasmic reticulum Ca2^++^ ATPase (SERCA) inhibitor Thapsigargin (TPG) for 2 h before harvesting and western blots were probed with the indicated antibodies. Blots from one representative experiment out of three are shown in the figure. (**B**) The intensity of the specific bands was quantified by densitometry in three independent experiments. The fold change relative to the untreated control is shown. Significance was calculated by Student t-test. * = P≤0.05. (**C**) Blockade of the ISR prevents the upregulation of ATF4 in BPLF1-expressing cells. BPLF1 transfected HeLa cells were cultured overnight and then treated with the indicated concentrations of the integrated stress response inhibitor (ISRIB) that reverses the effect of eIF2α phosphorylation, the PKR inhibitor CAS 608512-97-6 that prevents triggering of the ISR by foreign nucleic acids, the PERK inhibitor CAS1337531-89-1 that halts the ISR triggered by ER stress, and the GCN2 inhibitor CAS 1448693-69-3 that blocks the ISR triggered by tRNA depletion and ribosomal stress for 2 h before harvesting. (**D**) Quantification of the specific bands in four independent experiments. The data are shown as a ratio between the intensity of the FLAG-BPLF1 and ATF4 bands after normalization for the intensity of the GAPDH loading control. Significance was calculated by Student t-test. *** = P≤0.001; ** = P≤0.01

To further validate the involvement of eIF2α phosphorylation and investigate which eIF2α kinase mediates the effect, the upregulation of ATF4 was compared in BPLF1 transfected HeLa cells treated with the integrated stress response inhibitor (ISRIB) that counteracts the effect of eIF2α phosphorylation (Sidrauski et al., 2013), a PKR inhibitor that prevents triggering of the ISR by foreign nucleic acids (Jammi et al., 2003), a PERK inhibitor that halts the ISR triggered by ER stress (Axten et al., 2012), and a GCN2 inhibitor that blocks the ISR triggered by tRNA depletion and ribosome stalling (Brazeau and Rosse, 2014). The upregulation of ATF4 was significantly reduced by ISRIB and the GCN2 inhibitor, while the PKR and PERK inhibitors had minor and poorly reproducible effects (Fig 5C, 5D). Collectively, these findings point to the induction of persistent ribosomal stalling via functional inactivation of the ZNF598 ligase as the primary mechanism for induction of the ISR in cells expressing the vDUB.

### BPLF1 promotes the readthrough of stall-inducing mRNAs

Functional inactivation of ZNF598 by knockdown, mutation of the 40S ubiquitination sites, or overexpression of the USP21 deubiquitinase allows the progression of translation across stall-inducing mRNAs (Garshott et al., 2020; Sundaramoorthy et al., 2017). To investigate whether BPLF1 promotes translation through stall-inducing sequences, we utilized a dual fluorescence reporter system that expresses, from a single mRNA, GFP and RFP separated by linker encoding a FLAG-tagged villin headpiece (VHP) alone (K0) or fused to a stall-inducing stretch of twenty consecutive Lys (K20) (Shao et al., 2013) (Figure 6A). The linker regions are flanked by Ribosomal 2A skipping sequences (P2A), which allows independent assessment of translation before and after the linker. The K0 and K20 reporters were co-transfected in HEK293T cells with FLAG- BPLF1/BPLF1^C61A^, and the intensity of GFP and RFP was measured by FACS. A linear correlation of GFP and RFP fluorescence was observed in cells expressing the K0 reporter independently of expression of the catalytically active or mutant BPLF1 (Figure 6B upper panels). This allowed the gating of a readthrough region (R) where the transfected cells exhibited comparable levels of green and red fluorescence. Upon transfection of the K20 reporter, a robust repression of RFP fluorescence caused the accumulation of approximately 80% of the cells into a stalling region (S) where RFP fluorescence fell below the R threshold (Figure 6B lower panel left). A similar proportion of stalled cells was observed upon transfection of the catalytic mutant BPLF1^C61A^ (Figure 6B right panel), while expression of the active vDUB resulted in a very significant rescue of RFP fluorescence (Figure 6B middle panel). The results were highly reproducible, with more than 80% of the K20/BPLF1 transfected cells found in the R quadrant versus less than 20% in control and K20/BPLF1^C61A^ transfectants (Figure 6C). Despite the rescue, the RFP fluorescence remained lower in K20/BPLF1 transfected compared to K0/BPLF1 transfected cells (compare Figure 6B upper and lower panels). This was confirmed in western blots probed with antibodies to GFP and RFP, which revealed a significantly lower RFP:GFP ratio in K20/BPLF1 transfected compared to control cells (Figure 6D). This is likely explained by the capacity of BPLF1 to promote the readthrough of stall-inducing mRNAs in multiple reading frames, as shown to occur upon loss of ZNF598 (Sundaramoorthy et al., 2017). In line with this possibility, the expression of the FLAG-VHP-K20 fragment in BPLF1 transfected cells was comparable to that of FLAG-VHP-K0 in control cells (Figure 6D).

**Figure 6.**
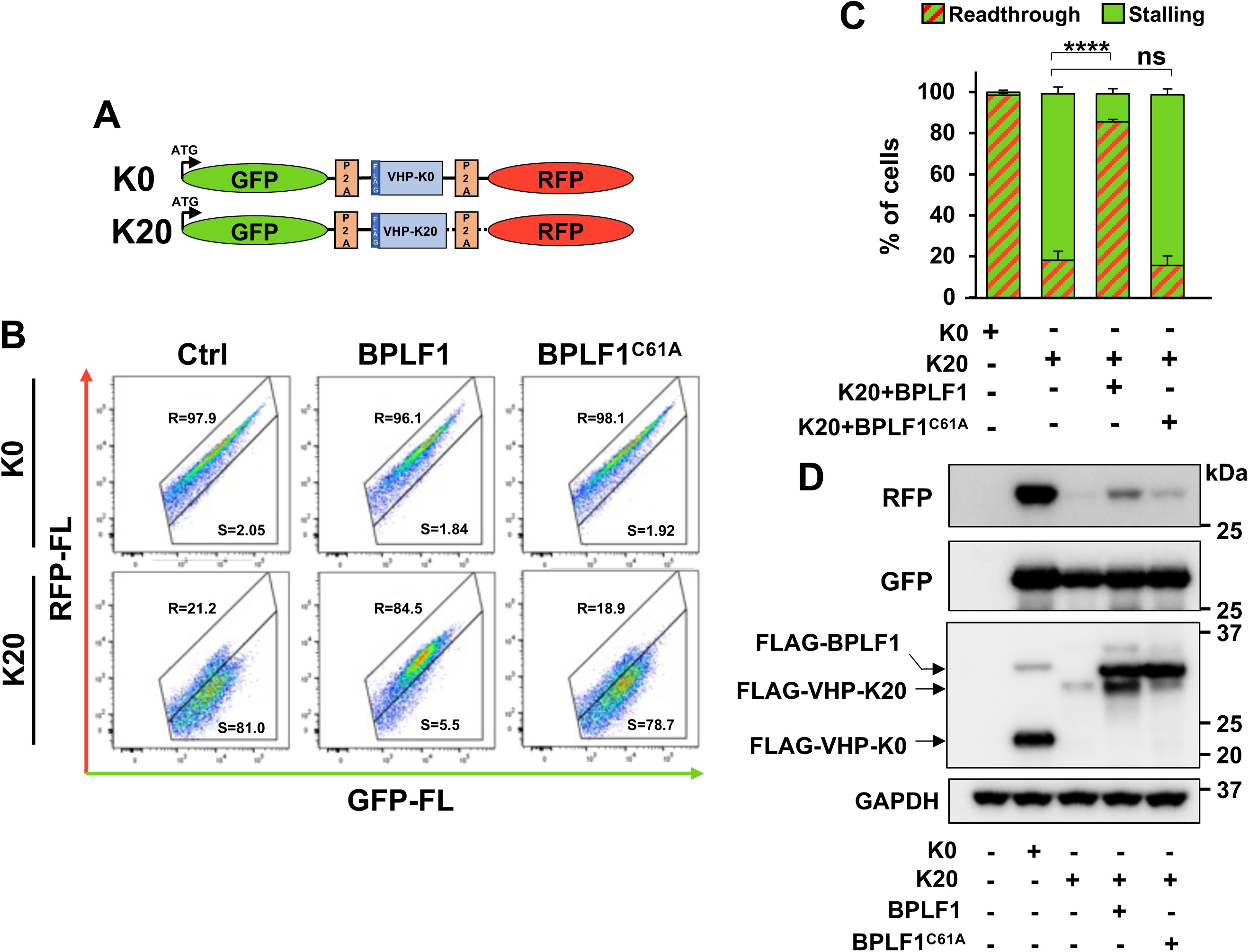
BPLF1 promotes translation elongation across a stall-inducing poly(A) sequence. (**A**) Schematic illustration of the translation-readthrough dual fluorescence reporter. The reporter expresses, from a single mRNA, the GFP and RFP separated by linker encoding a FLAG-tagged villin headpiece (VHP) alone (K0) or fused to a stall-inducing stretch of 20 consecutive lysins residues (K20). The linker regions are flanked by Ribosomal 2A skipping sequences (P2A), which allows independent assessment of translation before and after the linker. (**B**). BPLF1 promotes the readthrough of stall-inducing mRNAs. FLAG-BPLF1/BPLF1^C61A^ transfected HEK293T cells were co-transfected with either the K0 (upper panels) or the K20 reporter (lower panels) before analysis of GFP and RFP fluorescence by FACS. A readthrough (R) region was gated in the plots of K0 transfected cells exhibiting linear correlation between GFP and RFP fluorescence. Cells falling in the stalling (S) region exhibited decreased RFP:GFP fluorescence ratios. FACS plots from one representative experiment out of two are shown in the figure. (**C**). Quantification of the percentage of transfected cells falling in the (R) and (S) quadrants in two independent experiments. (**D**). BPLF1 promotes mRNA readthrough in multiple reading frames. Western blot of cell lysates from the experiment shown in Figure 5B were probed with the indicated antibodies. The GFP, RFP, and FLAG-VHP products were equally expressed in the K0 transfected cells. The RFP and FLAG-VHP-K20 products were virtually undetectable in control cells and cells expressing the BPLF1^C61A^ mutant. In BPLF1 transfected cells, GFP and FLAG-VHP- K20 were expressed at comparable levels, while RFP was significantly decreased.

### Endogenously expressed BPLF1 enhances the translation of viral proteins and virus release during productive EBV infection

Although reporters containing long poly(A) sequences have been instrumental in elucidating the mechanisms and consequences of ribosome stalling, their relevance for translation elongation is uncertain since protein coding open reding frames (ORFs) do not contain long poly(A) stretches. Thus, examples of how the system may operate when confronted with natural substrates are still scarce. ORF features that may affect translation elongation include the presence of Guanin-rich domains that may form high-order structures known as G-quadruplex (G4) (Song et al., 2016). The G4-forming sequences usually encode low-complexity amino acid repeats or short motifs frequently found in viral proteins. A prototype example is the ORF of the EBV latency maintenance protein EBNA1, which contains an approximately six hundred nucleotide long G-rich stretch encoding a Gly-Ala repeat (GAr) (Frappier, 2015) (Fig 7A). The GAr was shown to regulate the translation elongation of EBNA1 mRNA both *in vitro* and in cells, resulting in low protein expression and impaired antigen presentation (Murat et al., 2014).

**Figure 7.**
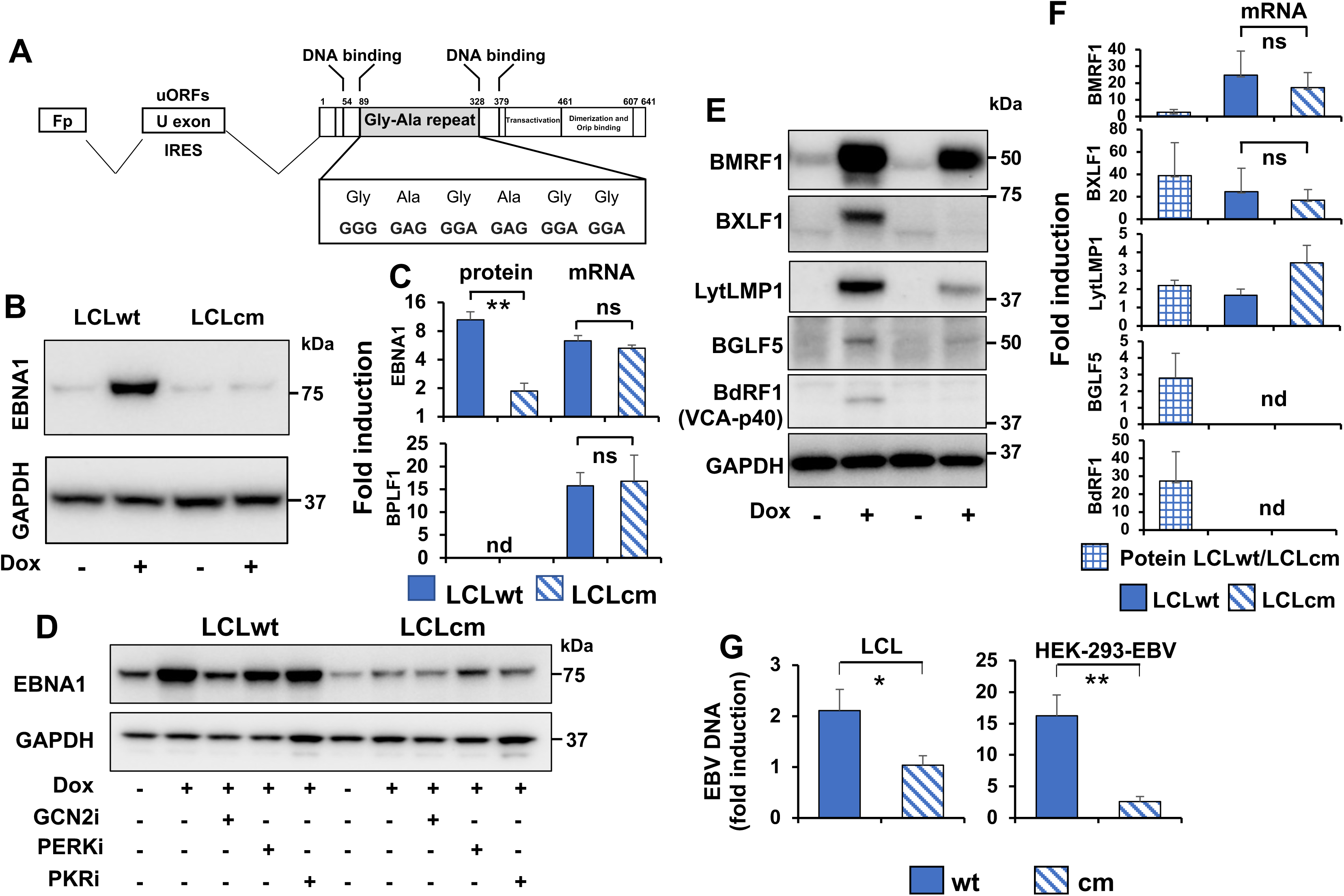
Catalytically active BPLF1 promots the translation of viral mRNAs during productive EBV infection and the release of infectious virus particles. (**A**) Schematic illustration of the EBNA1 mRNA transcriptional unit and coding sequence. During productive infection, the EBNA1 mRNA is transcribed from the Fp promoter, and exons located in the BamH-U and BamH-K fragments of the genome are joined by splicing. The short U exon contains IRES sequence and uORF domains. The protein coding ORF is in the K exon. The amino acid position of the DNA-binding, Gly-Ala repeat, transactivation, dimerization, and Orip binding domains are indicated. The amino acid and nucleotide sequences of a core GAGAGG motif repeated 13 times in the EBNA1 encoded by the prototype B95.8 virus are highlighted. (**B**) Catalytically active BPLF1 promotes the upregulation of EBNA1 during productive EBV infection. The productive virus cycle was induced by treatment with Dox for 72 h in LCLs immortalized with recombinant EBV expressing the wild-type (LCLwt) or catalytic mutant (LCLcm) BPLF1 and transduced with a Dox-regulated BZLF1 transactivator. Cell lysates were probed with the indicated specific antibodies. Blots from one representative experiment out of four are shown in the figure. (**C**) Quantification of the EBNA1 band detected in western blots and qPCR quantification of the corresponding mRNAs. Western blots for BPLF1 were not done due to a lack of antibodies detecting the endogenous protein but the comparable upregulation of the transcripts confirms similar induction levels. Fold induction was calculated relative to uninduced cells after normalization to GAPDH for western blots or the housekeeping MLN51 for qPCR. The mean ± SD of four independent experiments is shown. Significance was calculated by Student t-test. ** = P<0.01. (**D**) The upregulation of EBNA1 is dependent on GCN2 activity. The productive cycle was induced in LCLwt/cm in the presence of 1 μM of the GCN2 inhibitor, 1 μM of the PERK inhibitor or 0.3 mM of the PKR inhibitor. Blots from one representative experiment out of three are shown in the figure. (**E**). Catalytically active BPLF1 promotes the expression of a selection of early and late viral proteins. Western blots of cell lysates from the induction experiments shown in (B) were probed with the indicated antibodies. (**F**). Densitometric quantification of protein and mRNA expression in two to four independent experiments. Due to undetectable expression in uninduced cells, protein levels are expressed as the ratio of band intensity in cells expressing wild-type and mutant BPLF1. (**G**). The productive virus cycle was induced in HEK293T and LCL cells carrying EBV recombinant viruses expressing wild-type or catalytic mutant BPLF1 by transfection of a plasmid expressing the BZLF1 transactivator (HEK293T pair) or Dox treatment (LCL pair) and the amount of released enveloped virus was quantified in three days culture supernatants after treatment with DNase to remove free virus and DNA released by dead cells. The mean ± SD fold induction relative to control uninduced cells in three independent experiments for each cell pair is shown in the figure. Significance was calculated by Student t-test. ** = P<0.01.

To investigate whether physiological levels of the vDUB may regulate the translation of EBNA1 and other viral proteins, the productive virus cycle was induced in a pair of lymphoblastoid cell lines (LCLs) established by immortalization of B lymphocytes with recombinant EBV encoding either wild-type (LCLwt) or mutant BPLF1 (LCLcm). Since the virus is tightly latent in LCL cells, stable sublines expressing the Dox-regulated EBV transactivator BZLF1 were produced to ensure efficient and synchronous induction (Li et al., 2021). Control and Dox-treated cells were cultured for 72 h before protein and mRNA expression analysis by western blot and qPCR. We first tested whether BPLF1 may enhance the translation of EBNA1. A cartoon of the EBNA1 transcriptional unit and protein coding sequence in shown in Figure 7A. During the productive cycle, the mRNA is transcribed from the Fp promoter and splicing of the primary transcript joins a short exon located in the BamH1-U fragment of the viral genome to the first rightward ORF of the BamH1-K fragment (Brink et al., 2001). The U exon contains an IRES sequence (Isaksson et al., 2003) that partially overlaps with an uORFs (Bencun et al., 2018). A highly significant increase of EBNA1 protein levels was observed in western blots of cells expressing catalytically active BPLF1, while hardly any change occurred in cells expressing the mutant BPLF1^C61A^ (Figure 7B). A comparable increase of EBNA1 and BPLF1 mRNA was detected in the two cell lines, confirming equal levels of induction, and supporting the conclusion that the active vDUB promotes the translation of EBNA1 mRNA. BPLF1 protein levels were not analyzed due to lack of antibodies detecting the endogenously expressed proteins. The effect on EBNA1 translation was dependent on activation of the GCN2 kinase as confirmed by the virtually complete blockade of EBNA1 upregulation upon treatment with a GCN2 inhibitor, while inhibitors of PKR or PERK had only minor effects (Figure 7D). To test whether BPLF1 has a similar effect on other viral proteins, the western blots were probed with a panel of antibodies specific for LMP1, a truncated version of which is selectively upregulated during productive infection and is required for efficient virus release (Ahsan et al., 2005), the early proteins BMRF1 that encodes the viral DNA processivity factor, BXLF1 encoding the viral thymidine kinase, and BGLF5 encoding the viral alkaline exonuclease, and the late protein BdRF1 encoding the p40 subunit of the viral capsid antigen (VCA). Although the detection efficiency varied considerably due to the quality of the specific antibodies, in all cases, the expression was higher in cells expressing catalytically active BPLF1 despite comparable, or in the case of lytic LMP1, lower levels of the corresponding mRNAs (Figure 7E, 7F). In line with the inefficient expression of viral proteins required for various steps of the productive cycle, lower amounts of enveloped virus particles were recovered in the culture supernatants of cells expressing the catalytic mutant vDUB (Figure 7G). It is noteworthy that while significant, the effect of wtBPLF1 on virus release was relatively small in LCLs, in line with the tight latency observed in this cell type, but more robust in a previously described pair of HEK293T-EBV cell lines carrying wt and mutant BPLF1 (Gupta et al., 2018) where the productive virus cycle was efficiently induced by transfection of the BZLF1 transactivator.

## DISCUSSION

A growing body of evidence highlighting the importance of the RQC in resolving issues arising during the translation of stall-inducing mRNAs points to a critical role in viral infection. Yet, although understanding how viruses hijack this cellular control machinery could provide important clues on its function and regulation, the impact of viral infection on the RQC remains largely unexplored. Here, we have uncovered an unexpected role of the viral DUB encoded by EBV in reprograming the RQC to favor the translation of viral proteins while preventing the degradation of aberrant translation products that may activate antiviral responses.

Using a co-immunoprecipitation, mass spectrometry, and validation approach, we have found that the vDUB encoded in the N-terminal domain of the EBV large tegument protein BPLF1 is recruited to protein complexes that regulate mRNA biogenesis and translation and include the RQC ubiquitin ligases ZNF598 and LTN1, and the UFM1 ligase UFL1 that controls the ER-RQC pathway. It is noteworthy that although BPLF1 is a huge cytosolic protein of more than three thousand amino acids, the small N-terminal catalytic domain is released by caspase cleavage and can diffuse between the nucleus and cytoplasm (Gastaldello et al., 2013), which underlies its pleiotropic effects on the host cell reprogramming that enables productive viral infection. Our results point to the capacity of BPLF1 to counteract the ubiquitination of 40S ribosome subunits as a critical event in the viral remodeling of protein translation and cellular stress responses. In line with the notion that upon sensing ribosome collision ZNF598 initiates a process leading to the disruption of aberrant translation products and recycling of ribosome particles (Brandman et al., 2012; Filbeck et al., 2022; Garzia et al., 2017; Ikeuchi et al., 2019; Juszkiewicz and Hegde, 2017; Vind et al., 2020), we found that BPLF1 inhibits the LTN1-dependent proteasomal degradation of RQC substrates generated by the stalling of translation on poly(A) sequences. The series of events that links ribosome stalling to its downstream effects is not fully understood. In the currently accepted scenario, the ubiquitination of RPS10 and RPS20 by ZNF598 serves as the initiating events (Garshott et al., 2020), followed by the ubiquitination of RPS2 and RPS3 by ZNF598 and RNF10, which may work as a triage signal to regulate the turnover of ribosome subunits (Garzia et al., 2021). We found that BPLF1 counteracts the ubiquitination of both RPS10 and RPS3, mimicking the activity of cellular DUBs, such as USP21, OTUD3, and USP10 (Garshott et al., 2020; Garzia et al., 2021) that serve as physiological regulators of the pathway. ZNF598 was previously shown to be coopted by poxviruses to promote the accumulation of viral proteins (DiGiuseppe et al., 2018; Sundaramoorthy et al., 2021). Interestingly, while dependent on ubiquitination of the 40S subunits, the effect on poxvirus translation was still associated with inhibition of the cellular RQC (Sundaramoorthy et al., 2021) and may be at least partly explained by the need for poxviruses to reprogram the quality control machinery to allow the translation of mRNAs containing unusual poly(A) leaders (DiGiuseppe et al., 2018).

We unexpectedly found that the vDUB rescues ER-RQC substrates committed to lysosomal degradation following the UFMylation of RPL26 on ribosomes that stall while translating ER-targeted polypeptides. The capacity of BPLF1 to inhibit the UFMylation of both co-transfected and endogenous RPL26, together with the presence in the BPLF1 interactome of putative UFMylation substrates involved in ER-associated events, including the recognition, targeting, translocation, and glycosylation of secretory and membrane proteins, strongly support its involvement in the regulation of UFMylation-driven processes. Our initial hypothesis that, in addition to the known specificity for ubiquitin and Nedd8 conjugates (Gastaldello et al., 2010), the vDUB may also serve as a deUFMylase was conclusively refuted by the failure of catalytically active BPLF1 to hydrolyze UFM1 conjugates *in vitro* and in cells, pointing to the ligase as the primary target of the inhibitory effect. UFL1 is the only known cellular UFM1 ligase and has both nuclear and cytosolic substrates (Liu et al., 2020; Qin et al., 2019; Tatsumi et al., 2010; Zhang et al., 2015). Its recruitment to the ER is mediated by binding to DDRGK1 (Cai et al., 2015; Liu et al., 2017), which is also required for optimal ligase activity (Peter et al., 2022). Our finding that catalytically active BPLF1 inhibits RPL26 UFMylation and prevents the interaction of UFL1 with DDRGK1 links a ubiquitination step to the assembly of the ligase. Thus, while the molecular interactions that couple ribosome stalling to the activation of ER-RQC are currently unknown, our results point to the sensing of ubiquitinated stalled ribosomes as a possible triggering event. It is important to note that we cannot formally exclude that the dependency of UFL1 inactivation on the expression of catalytic active BPLF1 may be an artifact due to the failure of the BPLF1^C61A^ mutant to properly fold and interfere with the binding of UFL1 to DDRGK1. This seems, however, unlikely given the comparable, if not superior, interaction of the mutant with UFL1 and all other members of the BPLF1 interactome. Independently of the mechanism of ligase inactivation, we found that BPLF1 inhibits two important consequences of RPL26 UFMylation, the lysosome-dependent degradation of ER-RQC substrates and the activation of ER-phagy. The capacity to coordinately prevent the disruption of both RQC and ER-RQC substrates is likely to afford a critical advantage to the virus because the aberrant products of faulty translation constitute a significant source of antigenic peptides recognized by the antiviral responses (Dolan et al., 2011). Interestingly, the unexpected effect of the late protein BPLF1 on the processing of both proteasome- and autophagy-targeted precursors may at least partially explain the paradoxical finding that the abundantly expressed late viral antigens are poor targets of EBV antigen-specific T-lymphocytes while latent and immediate early antigens are efficiently recognized (Abbott et al., 2013).

Impairment of the RQC and ER-RQC was shown to independently trigger the ISR, via a direct effect of persistent ribosome stalling on the activation of GCN2 (Wu et al., 2020; Yan and Zaher, 2021) or via the activation of PERK in response to the accumulation of unprocessed ER substrates (Liang et al., 2020; Wang et al., 2020), respectively. In line with the inhibition of RQC and ER-RQC, we found that BPLF1 activates the ISR, as illustrated by the accumulation of phosphorylated eIF2α, upregulation of ATF4 and CHOP, and decrease of global protein synthesis in cells expressing the active enzyme. Significantly, the upregulation of ATF4 was inhibited by treatment with ISRIB, which reverses the effect of eIF2α phosphorylation, and with a specific inhibitor of GCN2, while minor effects were observed upon inhibition of PERK or PKR. The predominant involvement of GCN2 further corroborates the central role of the prolonged ribosome stalling associated with the functional inactivation of ZNF598 in the translation reprogramming induced by the vDUB. While the association of viral infection with ISR activation is well known, the role of ISR in infection is unclear and is expected to vary depending on the virus and type of infection. Like many other viruses, EBV inhibits the activation of PKR, which prevents a total shutoff of protein synthesis that could affect virus replication (Cesaro and Michiels, 2021). The preferential shutoff of cellular protein synthesis may then be achieved through the degradation of host cell mRNAs by the viral RNase BGLF5 (Daly et al., 2020). However, viral mRNAs are also targeted, and the selectivity mechanism remains unknown. Our findings suggest a scenario where the activation of ISR may be instrumental for reprograming the translation machinery in favor of viral mRNAs that often contain Internal Ribosome Entry Sites (IRES) and upstream open reading frames (uORFs). Untranslated uORFs have been identified in all classes of latent, immediate early, early, and late EBV transcripts (Bencun et al., 2018). Several were shown to downregulate protein expression in luciferase reporter assays confirming their potential role in the regulation of EBV translation, but their activity during productive infection was not explored. It is noteworthy that, while uORFs are commonly regarded as translation inhibitors, under stress conditions, uORFs enhance the translation of mRNAs encoding ATF4 and other cellular proteins that regulate the cell cycle and promote stress resilience (Andreev et al., 2015; Barbosa et al., 2013). Thus, the activation of ISR may afford multiple advantages to EBV by redirecting translation towards viral proteins and a restricted set of cellular proteins required for survival and efficient virus production.

Inhibition of the RQC facilitates translation elongation across stall-inducing poly(A) sequences (Garshott et al., 2020; Juszkiewicz and Hegde, 2017; Sundaramoorthy et al., 2017). Hence, if stalling goes undetected, the ribosome may restart translation after prolonged pausing, albeit with an increased risk of frame shifts and amino acid misincorporation (Buchan and Stansfield, 2007). In line with this notion, we found that the functional inactivation of ZNF598 induced by BPLF1 correlates with a virtually complete readthrough across the poly(A) sequence of the GFP-VHPK20-RFP reporter, resulting in a similar expression of the VHPK0 and VHPK20 fragment, while the relatively lower expression of RFP is probably explained by the imprecise restarting of translation downstream of the poly(A) tract. Most importantly, we found that under physiological conditions of expression, catalytically active BPLF1 is required to efficiently translate viral mRNAs in productively infected cells. The highly significant upregulation of EBNA1 is particularly remarkable. EBNA1 is the viral episome maintenance protein of EBV and is the only viral protein expressed in all types of infected normal and malignant cells (Frappier, 2015). While the key to EBV persistence during latency, EBNA1 plays important roles during productive infection (Sivachandran et al., 2012) when it is expressed from a dedicated lytic promoter that produces transcripts containing both IRES (Isaksson et al., 2003) and uORF (Bencun et al., 2018) features. In latency, the expression of EBNA1 is maintained at very low levels due to inefficient translation mediated by the elongation-hindering effect of the G4 forming sequence in the GAr coding domain (Murat et al., 2014). Our findings point to two possible mechanisms by which BPLF1 could enhance the translation of EBNA1 during productive infection. Via inhibition of the RQC, BPLF1 may allow a slowdown of translation required to resolve the G4 by the ribosome itself or associated RNA helicases. In addition, the activation of ISR may facilitate translation driven by the IRES and uORFs contained in the mRNA transcribed from the lytic promoter. Based on the small panel tested in this work, it seems likely that one or both mechanisms may enhance the translation of other viral proteins, which could explain the dependency on the vDUB activity for the efficient release infectious viral particles.

Collectively our findings outline a scenario where, by counteracting the activity of the ZNF598 ligase, with consequent inhibition of the RQC and ER-RQC, the vDUB may protect the infected cells from immune recognition while facilitating the handling of challenging viral mRNAs by the ribosome. This translational bias is amplified by the activation of the GCN2 kinase that contributes to the viral-induced shutoff of global protein synthesis while favoring the translation of mRNAs containing features such as uORFs and IRES that are frequently found in viral transcripts. Genome-wide analysis has revealed an unusually high density of putative G4 forming sequences in the repeat and regulatory regions of human herpesviruses (Biswas et al., 2016). Like EBNA1, LANA1, the genome maintenance protein of the other known human oncogenic herpesvirus Kaposi Sarcoma virus (KSHV), contains a G4 forming repeat that inhibits translation and antigen presentation (Dabral et al., 2020; Kwun et al., 2011). Functional vDUB domains have been identified in all BPLF1 homologs studied to date (Masucci, 2022). It remains to be seen whether the capacity to hijack the RQC and IRS pathways to promote virus production is a shared property of the enzymes encoded by this family of human pathogens.

## ACKNOWLEDGMENTS

We are grateful to Ron Kopito, Department of Biology, Stanford University, Stanford, CA; Masaaki Komatsu, Department of Biochemistry, Niigata University Graduate School of Medical and Dental Sciences, JP; Yogesh Kulathu and Joshua Peters MRC Protein Phosphorylation and Ubiquitination Unit, University of Dundee, UK; Jaap M. Middeldorp, VU University Medical Center, Amsterdam, NL, and Nico Dantuma, Department of Cell and Molecular Biology, Karolinska Institutet, Stockholm, SE, for providing reagents and technical advice. This investigation was supported by grants awarded by the Swedish Cancer Society (contract 211402Pj01H), the Swedish Research Council, and the Karolinska Institutet, Stockholm, SE. FAA was supported by a postdoctoral fellowship awarded by the Wenner-Gren Foundation, Stockholm, SE; FE was supported by and Erasmus training fellowship from the University of Coimbra, PT; SX was supported by doctoral fellowship awarded by the China Scholarship Council

## METHODS

### Reagents

For a complete list of reagents, kits, and commercially available or donated plasmids with source identifiers, see Table S2. For a list of primers used for cloning and qPCR analysis, see Table S3.

### Plasmid construction

To construct the GFP-nonSTOP reporter, the stop codon was removed from the GFP-C1 plasmid using PCR primers listed in Table S3. The product was religated using the NEbuilder HiFi DNA Assembly Master Mix according to the manufacturer’s instructions. The FLAG-Tag was removed from the eGFP-ER-K20-ChFP reporter plasmid using the Q5® Site-Directed Mutagenesis Kit according to the recommended protocol. A plasmid for prokaryotic expression of UFSP2 was constructed by extracting the USFP2 coding sequence from the pCMV3-UFSP2-FLAG plasmid by PCR, followed by in-frame insertion into the pET28a vector using the NEbuilder HiFi DNA Assembly Master Mix. A plasmid for procaryotic expression of the catalytic mutant BPLF1^C61A^ was produced by site-directed mutagenesis of the pET28a-His-BPLF1 plasmid.

### Tandem mass spectrometry and bioinformatics analysis

The mass spectrometry characterization of the BPLF1 interactome was previously reported (Gupta et al., 2018). For the current analysis, proteins detected by at least four unique spectral counts in the FLAG-BPLF1 and FLAG-BPLF1^C61A^ immunoprecipitates, and either absent or detected at a significantly lower level in duplicate samples of immunoprecipitates from cells transfected with the FLAG-empty vector were considered as positive hits. The bioinformatics resource Search Tool for the Retrieval of Interacting Genes (STRING) database v. 9.0 (Snel et al., 2000) and DAVID (Huang da et al., 2009) were used to identify the overrepresentation of genes in particular functional categories. Analysis of biological process (BP), molecular function (MF), and cellular component (CC) terms was performed using the ToppCluster (Kaimal et al., 2010) and Gene Ontology (Ashburner et al., 2000; The Gene Ontology, 2019) databases. Protein interaction network analysis was performed using the STRING database v. 9.0 (Snel et al., 2000) and the interaction network was visualized using Cytoscape (Shannon et al., 2003).

### Cell lines

The HeLa, HEK293T, U2OS, and HCT116 cell lines were cultured in Dulbecco’s minimal essential medium (DMEM) supplemented with 10% FBS and 10 μg/ml ciprofloxacin (complete medium) and grown in a 37°C incubator with 5% CO2. TetOn- BZLF1-EBV immortalized lymphoblastoid cell lines (LCLs) expressing catalytically active or inactive BPLF1 (Li et al., 2021) were cultured in RPMI1640 medium supplemented with 10% Tet-free FBS. The cells were transfected using either the lipofectamine 3000 or jetOPTIMUS® DNA transfection reagents according to the protocols recommended by the manufacturers. A stable HCT116 subline expressing the TetOn-mCherry-eGFP-RAMP4 ER-phagy reporter (HCT116-EATR) was produced by lentivirus transduction. Briefly, the TetOn-mCherry-eGFP-RAMP4, psPAX2, and pMD2.G plasmids were co-transfected into HEK293T cells using the JetOPTIMUS® kit. After culture overnight in Tet-free complete medium, the medium was refreshed, and the cells were cultured for an additional 48 h to allow virus production. The virus-containing culture supernatant was briefly centrifuged and passed through a 0.45 mm filter to removed cell debris before aliquoting and storing at -80°C until use. HCT116 were infected with virus-containing supernatants for 24 h in the presence of 8 μg/ml polybrene, followed by replacement of the infection medium with a fresh complete medium. Transduced cells were sorted twice by Fluorescence-activated cell sorting (FACS) based on ChFP and eGFP fluorescence, and stably expressing cells were cultured in Tet-free DMEM complete medium. U2OS-ufsp2 knock-out cells were generated by co-transfection with the px330-UFSP2 sgRNA1 and px330-UFSP2 sgRNA2 plasmids and a GFP-expressing plasmid. After two rounds of FACS sorting, the knock-out of ufsp2 was confirmed in western blots probed with an UFSP2-specific antibody.

### Induction of the productive virus cycle and detection of EBV transcripts and released virus

The productive virus cycle was induced by culturing the LCLwt and LCLcm cells for 72 h in the presence of 1.5 μg/ml Doxycycline. Total RNA was extracted with the Quick-RNA MiniPrep kit with in-column DNase treatment according to the recommended protocol. A second DNase treatment was performed to eliminate all traces of EBV DNA, followed by purification with an RNA Clean and Concentrator Kit. One μg total RNA was reverse transcribed using the SuperScript VILO cDNA Synthesis kit and quantified using a LightCycler 1.2 instrument (Roche Diagnostic) with the LC FastStart DNA master SYBR green I kit and specific primers (Supplementary Table S3). The housekeeping gene MLN51 was used as an internal control. All reactions were performed in duplicate. The relative fold change of gene expression was determined with the comparative cycle threshold (2^−ΔΔCT^) method. In the HEK293-EBV cell line (Li et al., 2021), the productive virus cycle was induced by transfection of a plasmid expressing the EBV BZLF1 transactivator, using Lipofectamine 3000. The release of infectious virus was monitored in culture supernatants cleared of cell debris by centrifugation of 5 min at 14000 rpm at 4 ^0^C and treated with 20 U/ml DNase I to remove free viral DNA. Viral DNA was isolated using the DNeasy Blood & Tissue Kit (Qiagen, Hilden, Germany) and quantitative PCR was performed with primers specific for a unique sequence in EBNA1 (Supplementary Table S3).

### Immunoblotting and co-immunoprecipitation

Cells were incubated for 30 min on ice in NP-40 lysis buffer (50 mM Tris-HCl pH 7.6, 150 mM NaCl, 5 mM MgCl2, 1 mM EDTA, 1 mM DTT, 1% Igepal, 10% glycerol) supplemented with protease inhibitor cocktail. After centrifugation at 14000 rpm for 15 min at 4°C, the protein concentration of the supernatants was measured with a protein assay kit. Equal amounts of lysates were fractionated in acrylamide Bis-Tris 4-12% gradient gel (Life Technologies Corporation, Carlsbad, USA). After transfer to PVDF membranes (Millipore Corporation, Billerica, MA, USA), the blots were blocked in Tris-buffered saline (TBS) containing 0.1% Tween-20 and 5% non-fat milk. The membranes were incubated with the primary antibodies diluted in blocking buffer for 1 h at room temperature or overnight at 4°C followed by washing and incubation for 1 h with the appropriate horseradish peroxidase-conjugated secondary antibodies. The immunocomplexes were visualized by enhanced chemiluminescence. For immunoprecipitation, cells were harvested 48 h after transfection and lysed in NP40 lysis buffer supplemented with inhibitors (protease/phosphatase inhibitor cocktail, 20 mM NEM and 20 mM iodoacetamide) for 30 min on ice. For immunoprecipitations under denaturing conditions, the cells were lysed for 10 min on ice in a small volume of lysis buffer supplemented with 1% SDS, followed by dilution to 0,1% SDS. For co-immunoprecipitation, the lysates were incubated with 50 μl of anti-FLAG or anti-Myc conjugates agarose affinity gel for 3 h at 4°C with rotation. For co-immunoprecipitation of endogenous UFL1 and ZNF598, the cell lysates were incubated for 4 h with specific antibodies, followed by capture of the immunocomplexes with protein-G coupled Sepharose beads. The beads were washed with lysis buffer, and the immunocomplexes were eluted by boiling in 2xNuPAGE Loading buffer supplemented with Sample Reducing agent. All images were acquired using a ChemiDoc Imaging system (Bio-Rad), and the intensity of target bands was quantified using the ImageLab software.

### Immunofluorescence

The cells were grown on cover slides in 6-well plates for 24 h before transfection. Semiconfluent monolayers were transfected as described and cultured for an additional 24 h. The cells were then fixed in 4% formaldehyde PBS buffer for 20 min, followed by permeabilization for 15 min in 0,1% Triton-X 100, blocking in PBS containing 4% bovine serum albumin (BSA) for 40 min, and incubation with the indicated dilutions of primary antibodies for 1 h at room temperature. After washing 3x5 min in PBS supplemented with 0.1% Tween 20 (PBST), the cells were incubated for another 1 h with the appropriate Alexa Fluor-conjugated secondary antibodies. The nuclei were stained with 2.5 μg/ml DAPI (Sigma, D9542) in PBS for 10 min, and the cover slides were mounted cell side down on object glasses with Mowiol (Calbiochem, 475904) containing 50 mg/ml 1,4-diazabicyclo[2.2.2]octane (Dabco; Sigma, D-2522) as an anti-fading agent. Images were acquired using a confocal fluorescence laser scanning Zeiss LSM900 microscope and analyzed using the Fiji ImageJ software.

### In vitro UFMylation assays

Recombinant His-BPLF1, His-UFSP2, His-UBA5, GST-UFC1, His-UFM1, and His- UFL1/Strep-DDRGK1 proteins were expressed in BL21 bacterial cells by overnight incubation at 16°C with 10 μM IPTG. The cells were lysed in Base buffer (25 mM Tris, 300 mM NaCl, 10% glycerol, 2 mM DTT, pH 8.0) supplemented with Complete protease-inhibitor cocktail by sonication with 5 cycles of 30s on-20s off pulses at 20 kHz on ice. Lysates were centrifuged at 13000 rpm for 20 min at 4°C. Proteins were purified using NiNTA- or Glutathione-agarose beads. Imidazole or glutathione were removed by dialysis against the base buffer. His-UFL1 complexed with Strep-DDRGK1 were then purified on a StrepTrap XT column. All proteins were purified by size-exclusion chromatography on a Superose 6HR 10/30 column on the base buffer, concentrated on Microcon-10 filters and stored on 25 mM Tris, 50 mM NaCl, 10% glycerol, 2 mM DTT, pH 7.5 at -80°C until use. UFMylation reactions were performed as previously described (Peter et al., 2022). Briefly, 0.25 μM His-UBA5, 5 μM GST-UFC1, 1 μM His- UFL1/Strep-DDRGK1, 0.5 μM of H3/H4 tetramer and 10 μM His-UFM1 were incubated on reaction buffer (50 mM HEPES, 10 mM MgCl2, 5 mM ATP, pH 7.4), in the absence or presence of BPLF1 or UFSP2, at 37°C for 90 min. The reactions were stopped by adding LDS-sample buffer supplemented with 100 mM DTT, and western blots were probed with the indicated antibodies.

### De-UFMylase assay

U2OS-UFSP2-KO cells were co-transfected with a reporter plasmid expressing a UFM1- GFP fusion protein and plasmids expressing FLAG-BPLF1/BPLF1^C61A^ or the cellular de-UFMylase. After overnight culture, the cells were lysed on RIPA buffer (50 mM Tris, 150 mM NaCl, 1% Ipegal, 0.5% deoxycholic acid, 0.1% SDS, pH 7.4) supplemented with complete protease inhibitors cocktail and 100 mM NEM. Lysates were cleared by centrifugation at 13, 000 rpm for 30 min at 4°C. Proteins on cell lysates were resolved by SDS-PAGE and transferred to PVDF membranes. Proteins were detected by immunoblotting with specific antibodies.

### RQC and ER-RQC reporter assays

The GFP-nonSTOP or ER-K20 reporters were co-transfected in HEK293T cells with either FLAG-BPLF1, FLAG-BPLF1^C61A^, or the empty FLAG-vector as control. After 24 h, the cells were lysed in NP-40 lysis buffer, and protein expression was assessed by western blot. Where indicated, 10μM or 100 nM Bafilomycin-A1 were added to the cultures 6 h before harvesting to inhibit proteasome- or lysosome-dependent degradation, respectively.

### ER Autophagy Tandem Reporter (EATR) assay

The expression of the EATR reporter was induced in HCT116-EATR cells grown on cover slides by treatment for 24 h with 2 μg/ml Doxycycline before FLAG-BPLF1 or FLAG-BPLF1^C61A^ transfection. After culture for additional 24 h, the cells culture medium was removed by repeated PBS washing, and the cells were starved by culture for 16 h in Earl’s balanced salt solution (EBSS) medium. The cells were then fixed and stained with the anti-FLAG antibody as described. Images were acquired using a Zeiss LSM900 confocal microscope, and the number of red dots in ≈45 BPLF1/BPLF1^C61A^ positive or negative cells was counted manually.

### Surface sensing of translation, SUnSET assay

HeLa cells were grown in 6-well plates and semiconfluent monolayers were transfected with FALG-BPLF1 or FLAG-BPLF1^C61A^ for 48 h. Ten μg/ml of Puromycin were added during the last 10 min before harvesting. As a positive control of translation inhibition, the cells were pretreated with 10 μM Cycloheximide (CHX) for 5 min before adding Puromycin. Cells were then washed with cold PBS and lysed in NP40 lysis buffer supplemented with a protease inhibitor cocktail. Western blots of lysates from an equal number of cells were probed with a Puromycin-specific antibody. For immunofluorescence, cells grown on cover slides were transfected and treated as described above. The cover slides were then processed for immunofluorescence staining as described. Images were acquired using a fluorescence microscope (Zeiss LSM900), and fluorescence intensity was quantified using the Fiji ImageJ software.

### Translation-readthrough assay

HEK293T cells were plated into 6-well plates the day before transfection. Semiconfluent monolayers were co-transfected with the pmGFP-P2A-K0-P2A-RFP (K0) or pmGFP- P2A-K(AAA)20-P2A-RFP (K20) together either with FLAG-BPLF1, FLAG- BPLF1^C61A^ or the empty FLAG-vector as transfection control. After culturing for 24 h, one aliquot of the cells was lysed with NP-40 lysis buffer for western blot analysis, and the expression of GFP and RFP was measured in the remaining cells by flow cytometry using a BD LSR II SORP apparatus and the data were analyzed with the FlowJo software.

### Statistical analysis

Plotting and statistical tests were carried out with data obtained in two or more independent experiments using the Microsoft Excel software. No assumptions about data normality were made, and a two-tailed Students t-test was used to determine statistical significance. Statistical significance is indicated in figures as ns *P* > 0.05, \**P* ≤0.05, \*\**P* ≤ 0.01, \*\*\**P* ≤0.001, \*\*\*\**P* ≤ 0.0001.

## Supporting Information

**Table S1.** Functional annotation of the BPLF1-interacting proteins involved in translation and RQC.

| Protein | MW [kDa] | ΣCoverage | Unique Peptides | ΣPSMs | Functional annotation |
| --- | --- | --- | --- | --- | --- |
| EIF2S1 | 36,1 | 15,6 | 4 | 28 | Eukaryotic translation initiation factor 2 subunit 1. Functions in the early steps of protein synthesis by forming a ternary complex with GTP and initiator tRNA. The complex binds to a 40S ribosomal subunit followed by mRNA binding to form the 43S pre-initiation complex. Is a key component of the integrated stress response (ISR). Phosphorylation by metabolic-stress sensing protein kinases (EIF2AK1/HRI, EIF2AK2/PKR, EIF2AK3/PERK and EIF2AK4/GCN2) converts EIF2S1/eIF-2-alpha in a global protein synthesis inhibitor, leading to attenuation of cap-dependent translation and preferential translation of ISR-specific mRNAs, such as the transcriptional activators ATF4. |
| EIF3E2 | 52,2 | 11,5 | 4 | 8 | Eukaryotic translation initiation factor 4E type 2. Binds the 7-methylguanosine-containing mRNA cap during the early steps of translation initiation acting as a repressor. In contrast to EIF4E, it is unable to bind eIF4G, suggesting that it may acts by blocking assembly of the IF4F complex. |
| EIF4G1 | 154,7 | 7,8 | 8 | 23 | Eukaryotic translation initiation factor 4 gamma 1; Component of the eIF4F complex involved in the recognition of the mRNA cap. Participates in the ATP-dependent unwinding of 5'-terminal secondary structure and recruitment of mRNA to the ribosome. |
| EIF4G2 | 98,1 | 4,1 | 3 | 5 | Eukaryotic translation initiation factor 4 gamma 2. May play a role in the switch from cap-dependent to IRES-mediated translation during mitosis, apoptosis, and viral infection. Cleaved by caspases and viral proteases. |
| EIF5A | 16,1 | 19,7 | 3 | 18 | Eukaryotic translation initiation factor 5A; mRNA-binding protein involved in translation elongation. Function in mRNA turnover, probably acting downstream of de-capping. Involved in actin dynamics and cell cycle progression, mRNA decay and maintenance of cell wall integrity. |
| KHSRP | 73,0 | 8,6 | 5 | 9 | Far upstream element-binding protein 2. Part of a ternary complex that binds to the downstream control sequence (DCS) of the pre-mRNA. Mediates exon inclusion in transcripts that are subject to tissue-specific alternative splicing. May interact with single-stranded DNA from the far-upstream element (FUSE). Also involved in the degradation of mRNAs that contain AU-rich elements (AREs) in their 3'-UTR. |
| RPLP2 | 11,7 | 38,3 | 3 | 10 | 60S ribosomal protein L2, mitochondrial |
| RPL8 | 22,4 | 22,4 | 3 | 9 | 60S ribosomal protein L8 |
| RPL9 | 20,9 | 36,8 | 6 | 35 | 60S ribosomal proteins 9, belongs to the L6P family of ribosomal proteins. |
| RPL13 | 18,8 | 20,1 | 5 | 43 | 60S ribosomal protein L13 |
| RPL13A | 23,6 | 20,7 | 4 | 39 | 60S ribosomal protein L13a. Associated with ribosomes but is not required for canonical ribosome function. Upon interferon-gamma induced phosphorylation dissociates from the ribosome and assembles into the GAIT complex which binds to stem loop-containing GAIT elements in the 3'-UTR of inflammatory mRNAs. |
| RPL15 | 20,5 | 19,5 | 3 | 27 | 60S ribosomal protein L15 |
| RPL17 | 19,6 | 34,3 | 5 | 40 | 60S ribosomal protein L17. Belongs to the ribosomal protein uL22 family |
| RPL18A | 16,7 | 33,3 | 5 | 32 | 60S ribosomal protein L18A. Belongs to the ribosomal protein eL20 family. |
| RPL26 | 11,6 | 19,6 | 4 | 28 | 60S ribosomal protein 26. Belongs to the ribosomal protein L24P family Is located near the docking sites for the translocation machinery signal recognition particle (SRP), the SEC61 translocon, and the oligosyl transferase (OST) complex. |
| RPL30 | 12,6 | 35,1 | 3 | 17 | 60S ribosomal protein L30 |
| RPL34 | 13,3 | 16,2 | 3 | 4 | 60S ribosomal protein L34 |
| RPL36 | 12,2 | 21,9 | 3 | 14 | 60S ribosomal protein L36 |
| RPL38 | 8,2 | 50,0 | 4 | 24 | Ribosomal protein L38; Belongs to the ribosomal protein eL38 family |
| RPS11 | 18,4 | 32,3 | 4 | 10 | Ribosomal protein S11; Belongs to the ribosomal protein uS17 family |
| RPS19 | 16,1 | 23,5 | 5 | 38 | 40S ribosomal protein S19 |
| RPS5 | 14,8 | 17,2 | 3 | 18 | 40S ribosomal protein S5, small ribosomal subunit uS7 |
| RPS7 | 22,1 | 25,8 | 5 | 27 | 40S ribosomal protein S7; Required for rRNA maturation; Belongs to the ribosomal protein eS7 family |
| RPN1 | 68,5 | 26,5 | 12 | 35 | Dolichyl-diphosphooligosaccharide--protein glycosyltransferase subunit 1; Essential subunit of the N-oligosaccharyl transferase (OST) complex that catalyzes the transfer of a high mannose oligosaccharide from a lipid-linked oligosaccharide donor to an asparagine residue within an Asn-X-Ser/Thr consensus motif in nascent polypeptide chains. The complex binds to ribosome to take part in co-translational protein folding. Following UFMylation of the N-terminal cytoplasmic domain, RPN1 delivers a stress ER stress signal during peptide translocation, which can ultimately induce ER-phagy or UPR. The protein forms part of the regulatory subunit of the 26S proteasome and may mediate binding of ubiquitin-like domains to the proteasome. |
| SEC63 | 87,9 | 7,6 | 4 | 11 | Translocation protein 63 homolog; ER membrane protein, member of the Sec61 complex that mediates co-translational the transport of polypeptides across the ER. Cooperates with SEC62 to facilitate targeting to into the SEC61 channel-forming translocon. The complex may also perform retrograde transport of ER proteins that are subject to the ubiquitin-proteasome-dependent degradation. |
| SRP68 | 66,1 | 10,5 | 6 | 19 | Signal recognition particle (SRP) subunit 68. The SRP targets secretory proteins to the rough ER. SRP68 binds the 7S RNA, which recruits SRP72 to the complex. The ribonucleoprotein complex may interact with docking proteins in the ER membrane and possibly participate in elongation arrest. |
| SRPRA | 66,5 | 7,2 | 4 | 20 | Signal recognition particle receptor subunit alpha. Component of the SRP receptor complex. In conjunction with the SRP complex ensures the correct targeting of the nascent secretory proteins to the ER membrane. Forms a guanosine 5'-triphosphate (GTP)-dependent complex with the SRP subunit SRP54. |
| SRPRB | 29,7 | 46,1 | 9 | 23 | Signal recognition particle receptor subunit beta; Component of the SRPR Has GTPase activity and may mediate the membrane association of SRPR. |
| UFL1 | 82,0 | 8,9 | 4 | 7 | UFM1 E3 ligase. Is recruited to the ER membrane by DDRGK1 and mediates UFMylation of RPN1 and RPL26/uL24, thereby promoting ER-phagy, which inhibits the unfolded protein response (UPR). DDRGK1 and CDK5RAP3 serve as substrate adapters for UFMylation and ER-phagy. Catalyzes UFMylation of various subunits of the ribosomal complex or associated components, such as RPS3/uS3, RPS20/uS10, RPL10/uL16, RPL26/uL24 and EIF6. |

**Table S2.** Reagents used in this paper.

| Reagent | Source | Identifier |
| --- | --- | --- |
| **Antibodies** |  |  |
| Rabbit polyclonal anti-Histone H3 | Millipore | Cat# 07-690 |
| Rabbit monoclonal anti-UFSP2 | Abcam | Cat# ab185965 |
| Mouse monoclonal anti-β-actin clone AC-15 | Sigma-Aldrich | Cat# A5441, RRID: AB_476744 |
| Mouse monoclonal anti-GAPDH | Millipore | Cat#CB1001 |
| Mouse monoclonal anti-tubulin | Millipore | Cat#CP06 |
| Mouse monoclonal anti-FLAG | Sigma-Aldrich | Cat# F3165, RRID: AB_259529) |
| Rabbit polyclone anti-FLAG | Sigma-Aldrich | Cat#F7425, RRID: AB_439687 |
| Anti-FLAG M2 affinity gel | Sigma-Aldrich | Cat#A2220 RRID:AB_10063035 |
| Rabbit monoclonal anti-Myc Tag | Cell signaling technology | Cat#2278 |
| Mouse monoclonal anti-S Tag, | Millipore | Cat#71549-3 RRID: AB_11210600 |
| Mouse anti-Myc Tag agarose (4A6) | Millipore | Cat# 16-219 |
| S-Protein Agarose | Novagen | Cat#69704 |
| protein-G coupled Sepharose beads | GE Healthcare | Cat#17-0885-01 |
| GammaBind Plus Sepharose | Cytiva | Cat#17088601 |
| Rabbit IgG isotype control | Abcam | Cat# ab172730 RRID:AB_2687931 |
| Donkey anti-Mouse IgG (H+L), Alexa Fluor 555 | Thermo Fisher Scientific | Cat# A-31570, RRID: AB_2536180 |
| Donkey Anti-Rabbit IgG (H+L) Antibody, Alexa Fluor 488 | Thermo Fisher Scientific | Cat# A-21206, RRID: AB_2535792 |
| Goat Anti-Rabbit IgG (H+L) Antibody, Alexa Fluor 647 | Thermo Fisher Scientific | Cat#A21245 RRID: AB_141775 |
| Rabbit polyclonal anti-ZNF598 | Invitrogen | Cat# PA559777 RRID: AB_2650226 |
| Rabbit polyclonal anti-LTN1 | GeneTex | Cat# GTX18154 |
| Rabbit polyclonal anti-UFL1 | Bethyl Laboratories | Cat# A303-456A, RRID: AB_10951658 |
| Rabbit polyclonal anti-DDRGK1 | Thermo Fisher Scientific | Cat# PA5-100467 RRID: AB_2849977 |
| Rabbit monoclonal anti-UFC1 | Abcam | Cat#ab189251 |
| Rabbit Monoclonal anti-UFM1 | Abcam | Cat# ab109305, RRID: AB_10864675 |
| Rabbit monoclonal anti-UFSP2 | Abcam | Cat# ab185965 |
| Rabbit polyclonal anti-SEC63 | Bethyl Laboratories | Cat# A305-084A, RRID: AB_2631479 |
| Rabbit polyclonal anti-RPN1 | Bethyl Laboratories | Cat# A305-026A, RRID: AB_2621220 |
| Mouse monoclonal anti-RPN1 | Santa Cruz Biotechnology | Cat# sc-48367, RRID: AB_628221 |
| Rabbit polyclonal anti-SRPRB | Bethyl Laboratories | Cat# A305-440A,<br>RRID: AB_2631831 |
| Rabbit polyclonal anti-SRP68 | Bethyl Laboratories | Cat# A303-955A,<br>RRID: AB_2620304 |
| Mouse monoclonal anti-VCP | Abcam | Cat# ab11433<br>RRID:AB_298039 |
| Rabbit monoclonal anti-RPS10 | Abcam | Cat# ab151550<br>RRID:AB_2714147 |
| Rabbit monoclonal anti-RPS3 | Abcam | Cat# ab128995<br>RRID:AB_11145466 |
| Rabbit Polyclonal anti-RPL26 | Abcam | Cat# ab59567,<br>RRID: AB_945306 |
| Rabbit polyclonal anti-SEC61B | Thermo Fisher Scientific | Cat# PA3-015<br>RRID: AB_2239072 |
| Rabbit polyclonal anti-eIF4G2 (DAP5) | Bethyl Laboratories | Cat# A302-249A,<br>RRID: AB_1730977 |
| Mouse monoclonal anti-EIF5A | BD Biosciences | Cat# 611976<br>RRID:AB_399398 |
| Mouse monoclonal anti-eIF4G1 (2A9) | Abnova | Cat# H00001981-M10,<br>RRID: AB_606171 |
| Rabbit anti-Phospho-eIF2α-Ser51 (D9G8) XP | Cell Signaling Technology | Cat# 3398,<br>RRID: AB_2096481 |
| Mouse monoclonal anti-eIF2α (L57A5) | Cell Signaling Technology | Cat#2103,<br>RRID: AB_836874 |
| Rabbit monoclonal anti-ATF4 (D4B8) | Cell Signaling Technology | Cat# 11815,<br>RRID: AB_2616025 |
| Mouse monoclonal anti-CHOP (L63F7) | Cell Signaling Technology | Cat# 2895,<br>RRID: AB_2089254 |
| Mouse monoclonal anti-Puromycin (12D10) | Millipore | Cat# MABE343<br>RRID: AB_2566826 |
| Rat monoclonal anti-EBV-BPLF1 | MAB core facility, Helmholtz Center, Munich, Germany (van Gent et al., 2014) | N/A |
| Rat monoclonal anti-EBV-EBNA1 | MAB core facility, Helmholtz Center, Munich, Germany | N/A |
| Rabbit polyclonal anti-EBV-BdRF1 | Dr. Jaap M. Middeldorp, VU University Medical Center,Amsterdam, NL | N/A |
| Rabbit polyclonal anti-EBV-BGLF1 | Dr. Jaap M. Middeldorp, VU University Medical Center,Amsterdam, NL | N/A |
| Mouse monoclonal anti LMP1 (CS1-4) | Dako, Glostrup, Denmark | M0897 |
| Rabbit polyclonal anti-EBV BXLF1 | ASLA Biotech, Riga Latvia | N/A |
| Mouse monoclonal anti-EBV-BMRF1 | Dr. Jaap M.<br>Middeldorp, VU<br>University Medical<br>Center, Amsterdam, NL | N/A |
| **Chemicals, peptides, and recombinant proteins** |  |  |
| IGEPAL CA-630 | Sigma-Aldrich | I3021; CAS: 9002-93-1 |
| MgCl<sub>2</sub> | Merck | Cat# M1028 |
| Sodium dodecyl sulphate | Sigma-Aldrich | L3771; CAS:151-21-3 |
| N-Ethylmaleimide | Sigma-Aldrich | E1271; CAS:128-53-0 |
| Iodoacetamide | Sigma-Aldrich | I1149; CAS:144-48-9 |
| Sodium deoxycholate monohydrate | Sigma-Aldrich | D5670; CAS:145224-92-6 |
| Triton X-100 | Sigma-Aldrich | T9284; CAS:9002-93-1 |
| Bovine serum albumin | Sigma-Aldrich | A7906; CAS:9048-46-8 |
| Tween-20 | Sigma-Aldrich | P9416; CAS: 9005-64-5 |
| Trizma base | Sigma-Aldrich | 93349; CAS:77-86-1 |
| Ciprofloxacin | Sigma-Aldrich | Cat#17850; CAS:<br>85721-33-1 |
| Doxycycline cyclate | Sigma-Aldrich | D9891; CAS: 24390-14-5 |
| MG132 | Sigma-Aldrich | M7449; CAS:<br>200-664-3 |
| Anisomycin | Sigma-Aldrich | A5862; CAS:22862-76-6 |
| Cycloheximide | Sigma-Aldrich | C4859; CAS:66-81-9 |
| Bafilomycin A1 | Sigma-Aldrich | B1793; CAS:88899-55-2 |
| Thapsigargin | Sigma-Aldrich | T9033; CAS:67526-95-8 |
| Imidazole | Sigma-Aldrich | I5513; CAS: 288-32-4 |
| Polybrene | Sigma-Aldrich | TR-1003-G; |
| Complete protease inhibitors cocktail | Roche Diagnostic | Cat#04693116001 |
| Phosphatase inhibitor cocktail | Roche Diagnostic | Cat#04906837001 |
| DAPI | Sigma-Aldrich | Cat#9542<br>CAS:28718-90-3 |
| Mowiol | Calbiochem | Cat# 475904<br>CAS:9002-89-5 |
| HA-Ubiquitin-VS | R&D systems | Cat# U-212-025 |
| Histone H3.1/H4 Tetramer | NEB Biolabs | Cat# M2509S |
| Glutathione | ThermoFisher Scientific | Cat# 78259 |
| ATP | Thermo Fisher Scientific | Cat# R0441 |
| PERK inhibitor I, GSK2606414 | Sigma-Aldrich | 516535<br>CAS 1337531-89-1 |
| GCN2 inhibitor (A92) | AXON Medchem | Axon 2720;<br>CAS:1448693-69-3 |
| PKR inhibitor | Sigma-Aldrich | Cat# 527451<br>CAS 608512-97-6 |
| ISRIB | Sigma-Aldrich | SML0843<br>CAS:1597403-47-8 |
| **Kits** |  |  |
| DC Protein Assay quantification kit | Bio-Rad | Cat#A500-0116 |
| SuperSignal™ WestPico PLUS Chemiluminescent Substrate | Thermo Scientific | Cat#XE356732 |
| Lipofectamine 3000 transfection kit | Invitrogen | Cat#L3000-015 |
| jetOPTIMUS DNA transfection reagent | Polyplus | Cat#101000006 |
| Q5® Site-Directed Mutagenesis Kit | New England Biolabs | Cat#E0554S |
| NEBuilder® HiFi DNA Assembly Master Mix | New England Biolabs | Cat#E2621S |
| Quick-RNA MiniPrep kit | Zymo Research | Cat#R1055 |
| RNA clean and concentrator | Zymo Research | Cat#R1013 |
| SuperScript VILO cDNA Synthesis kit | Invitrogen | Cat#11754050 |
| LC FS DNA Master SYBR Green I | Roche | Cat#12239264001 |
| **Protein purification columns** |  |  |
| HisPur NiNTA resin | Thermo Scientific | Cat# 88222 |
| Pierce Glutathione Agarose | Thermo Scientific | Cat# 16101 |
| StrepTrap XT | Cytiva | Cat# 29401317 |
| Superose 6HR 10/30 | Cytiva | Cat# 17-0537-01 |
| Microcon-10 Centrifugal filter | Merck | Cat# MRCPR010 |
| **Experimental models: Cell lines** |  |  |
| RPMI1640 | Sigma-Aldrich | Cat# R8758 |
| DMEM | Sigma-Aldrich | Cat# D6429 |
| EBSS | Gibco | Cat# 1854705 |
| HeLa | ATCC | RR-B51S |
| HEK293T | ATCC | CRL3216 |
| U2OS | ATCC | HTB-96 |
| U2OS ufsp2 knock out | This paper | N/A |
| HCT116 | ATCC | CCL-247 |
| HCT116-EATR | This paper | N/A |
| HEK293T-EBV- BPLF1wt/cm | (Gupta et al., 2018) |  |
| LCLwt/cm (Tet-on BZLF1) | (Li et al., 2021) | N/A |
| **Recombinant DNA** |  |  |
| pCMV3-RPL26-Myc | Sino Biological | HG16834-CM |
| pCDNA3.1-RPL26-S | gift from Ron R. Kopito | N/A |
| pCMV3-UFSP2-FLAG | Sino Biological | HG17133-CF |
| pTRIPZ lentiviral vector | ThermoScientific | Cat#RHS4750 |
| pCW57.1 | gift from David Root | Addgene #41393 |
| psPAX2 | gift from Didier Trono | Addgene # 12260 |
| pMD2.G | gift from Didier Trono | Addgene # 12259 |
| TetOn-mCherry-eGFP-RAMP4 | gift from Jacob Corn | Addgene #109014 |
| pLenti-X1-Neo-DDRGK1-WT-HA | gift from Jacob Corn | Addgene # 139845 |
| eGFP-ER-K20-RFP | gift from Yihong Ye | Addgene # 133861 |
| pmGFP-P2A-K0-P2A-RFP | gift from Ramanujan Hegde | Addgene # 105686 |
| pmGFP-P2A-K(AAA)20-P2A-RFP | gift from Ramanujan Hegde | Addgene # 105688 |
| FLAG-HA-USP21 | gift from Wade Harper | Addgene # 22574 |
| px330-UFSP2 sgRNA1 | gift from Yihong Ye | Addgene # 134638 |
| px330-UFSP2 sgRNA2 | gift from Yihong Ye | Addgene # 134637 |
| pcDNA3-MYC-UFM1ΔC2 | Gift from Masaaki Komatsu | N/A |
| pcDNA1-UFM1-GFP | Gift from Yogesh Kulathu | N/A |
| pET15b-His-UBA5 | Gift from Yogesh Kulathu | N/A |
| pGX-6P1-GST-UFC1 | Gift from Yogesh Kulathu | N/A |
| pET-15b-His-UFM1 | Gift from Yogesh Kulathu | N/A |
| pETDuet1-His-UFL1-Strep-DDRGK1 | Gift from Yogesh Kulathu |  |

**Table S3.**
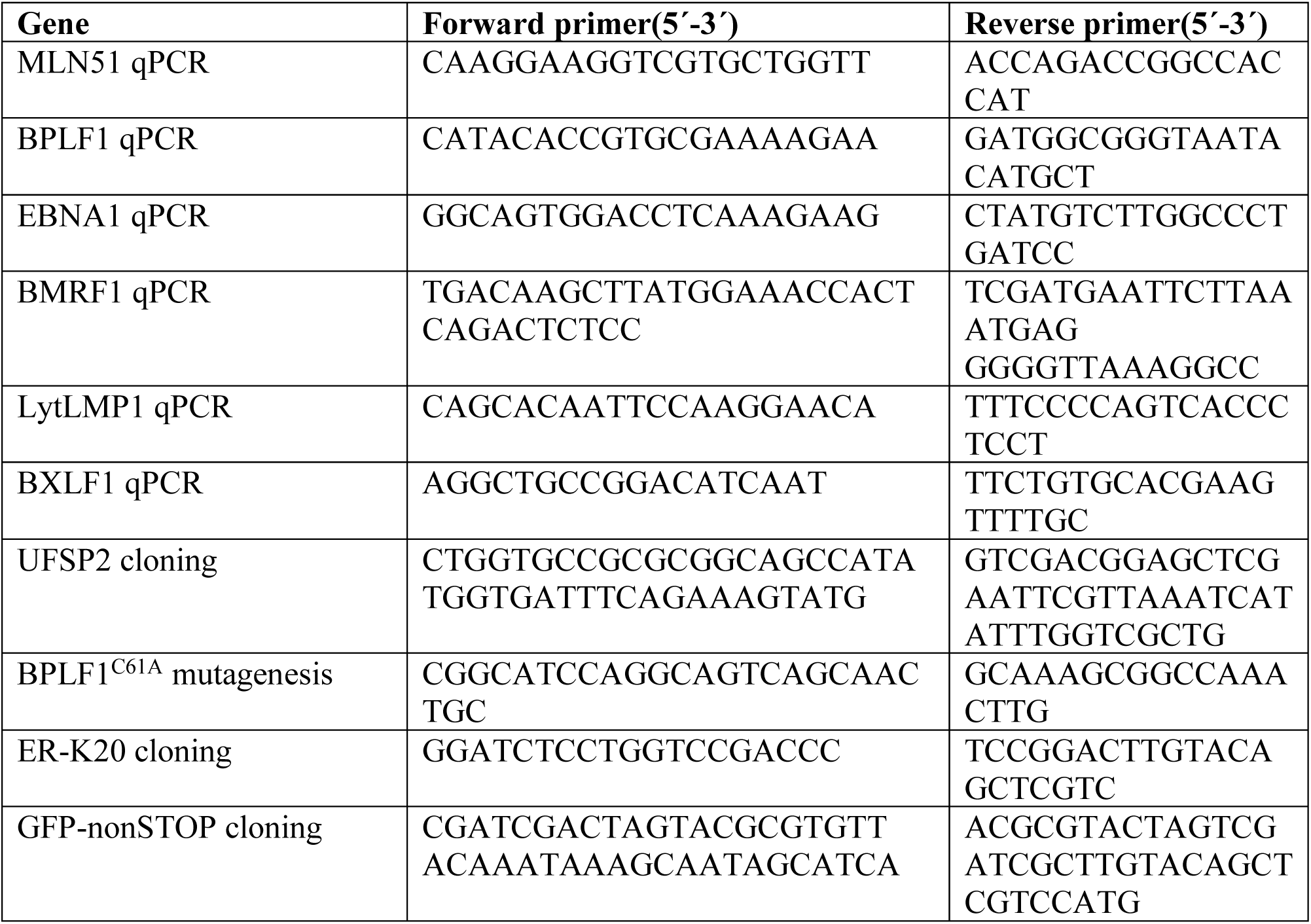
PCR and qPCR primers.

**Figure S1.**
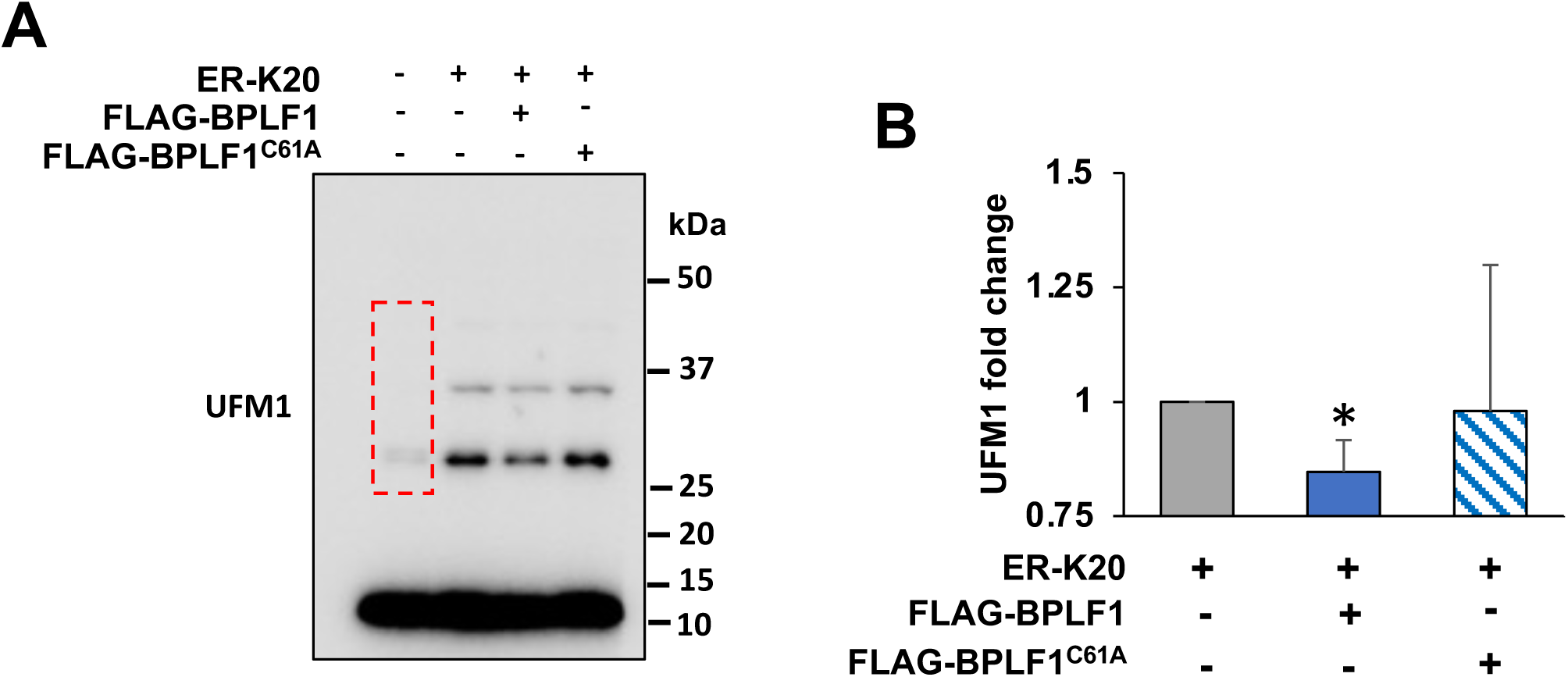
BPLF1 inhibits the UFMylation of endogenous RPL26 in HeLa cells expressing a ribosome stall-inducing reporter. (**A)** BPLF1 inhibits the UFMylation of endogenous RPL26 in cells expressing a ribosome stall-inducing reporter. HEK293T cells were co-transfected with FLAG-BPLF1/BPLF1^C61A^ and the stall-inducing ER-K20 reporter and western blots of cells harvested after 24h were probed with antibodies to UFM1. Western blots from one representative experiment out of three are shown in the figure. (**B**) Densitometry quantification of the UFMylated proteins (red dotted area in Fig S1A) in three independent experiments. Fold change was calculated relative to vector transfected cells. Statistical analysis was performed using Student t-test. *P ≤ 0.05

**Figure S2.**
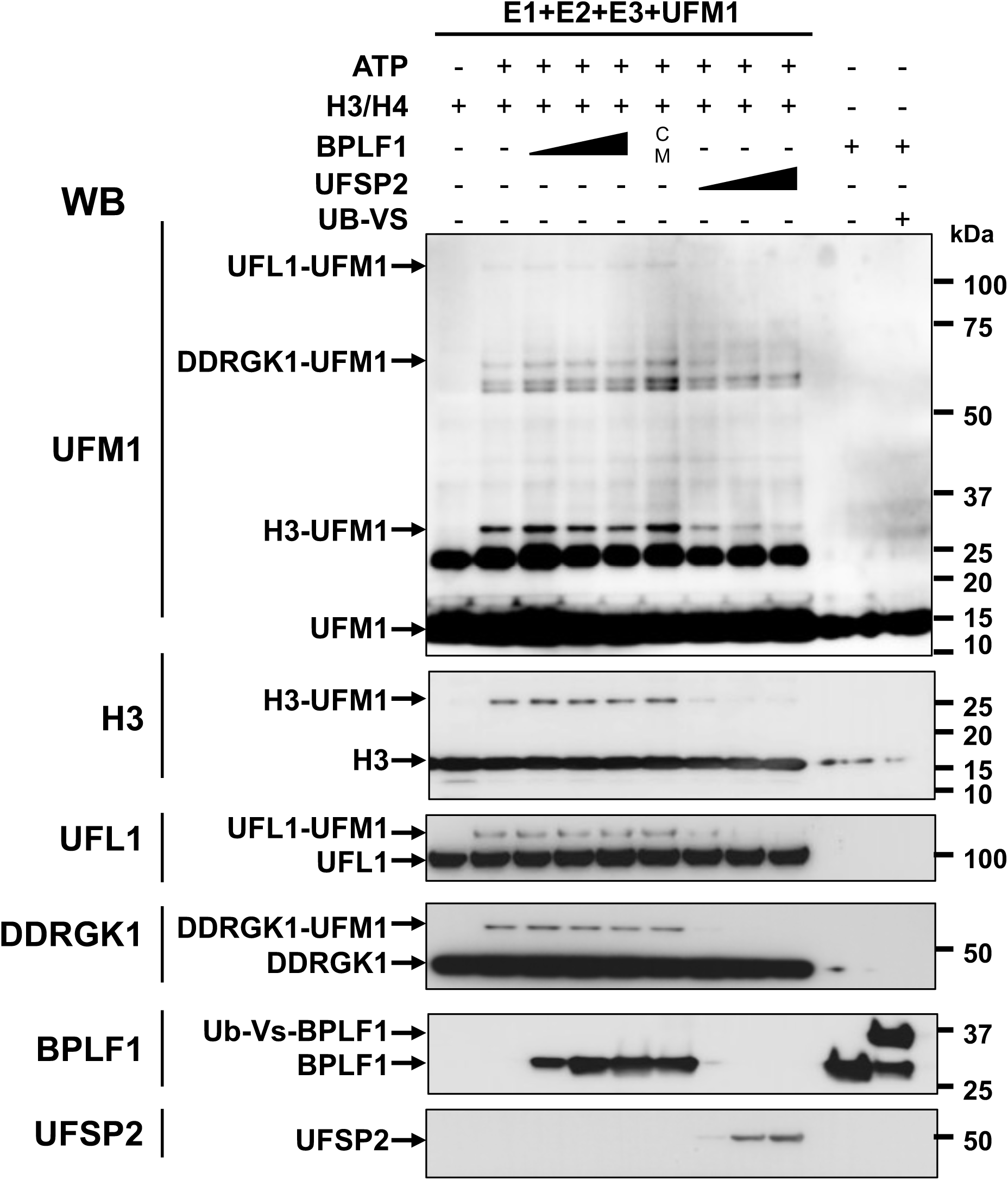
BPLF1 does not have de-UFMylase activity. Representative western blot of *in vitro* UFMylation reactions performed in the presence or absence of BPLF1 or UFSP2. UFMylation reactions were performed in the presence of purified components of conjugation cascade: 0.25 μM recombinant His-UBA5 (E1), 5 μM GST-UFC1 (E2), 1 μM His- UFL1/Strep-DDRGK1 (E3), 0.5 μM of H3/H4 complex (substrate), and 10 μM His-UFM1 in the absence (Lane 1) or presence (Lanes 2-9) of 5 mM ATP. Where indicated increasing concentrations of recombinant BPLF1 (lanes 3-5; 0.3, 0.75, and 1.5 μM, respectively), His-BPLF1^C61A^ (Lane 6; 1.5 μM) or His-UFSP2 (lane 7-9; 0.03, 0.075, and 0.15 μM, respectively). The DUB activity of recombinant BPLF1 was confirmed by labeling 1.5 μM His-BPLF1 with 1 μM of HA-Ubiquitin-Vinyl-sulphone (Ub-VS) (lane 10-11). All reactions were incubated at 37°C for 90 min and analyzed by western blot using specific antibodies. UFMylated H3, ULF1 and DDRGK1 were detected when the rection was performed in the presence of ATP (lane 2). The addition of increasing amounts of BPLF1 or BPLF^C61A^ had no effect (lane 3-6) while efficient de-UFMylation was induced by minute amounts of UFSP2 (lane 7-9).

**Figure S3.**
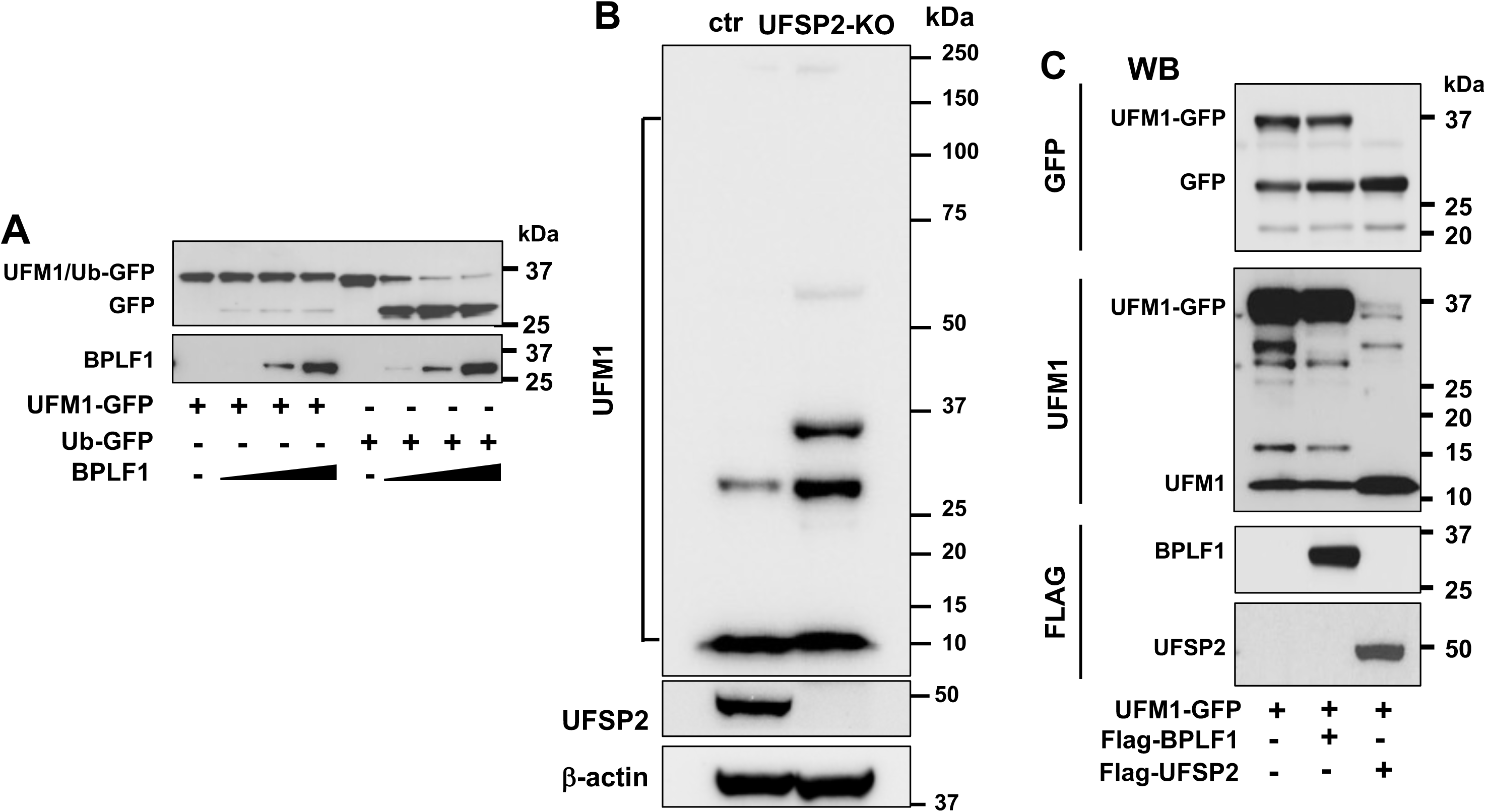
BPLF1 does not promote UFM1 deconjugation in cells. (**A**) BPLF1 does not cleave the UFM1-GFP reporter. Increasing amounts of purified recombinant BPLF1 (0.3, 0.75, 1.5 mM) were mixed with the UFM1-GFP or Ub-GFP reporters (0.3 mM) in reaction buffer and incubated for 1 h at 37°C followed by western blot analysis. One representative experiment out of three is shown in the figure. (**B**) Characterization of the UFSP2-KO cell line. The main cellular de-UFMylase UFSP2 was knocked-down in U2OS cells by co-transfection with GFP and px330-USP2-sgRNAs expressing plasmids. After two rounds of FACS sorting the expression of UFSP2 was assessed in western blots. A representative western blot illustrating the loss of UFSP2 and concomitant accumulation of endogenous UFMylated substrates is shown in the figure. (**C**) BPLF1 does not promote deUFMylation in cells. The USFP2-KO cells were co-transfected with an UFM1-GFP reporter plasmids and plasmids expressing either BPLF1 or the cellular de-UFMylase UFSP2. The production of free GFP and UFM1 was monitored in western blot probes with the specific antibodies. Representative western blots illustrating the failure of BPLF1 to cleave the reporter, whereas free GFP and UFM1 accumulated in cells expressing UFSP2. (**C**)

**Figure S4.**
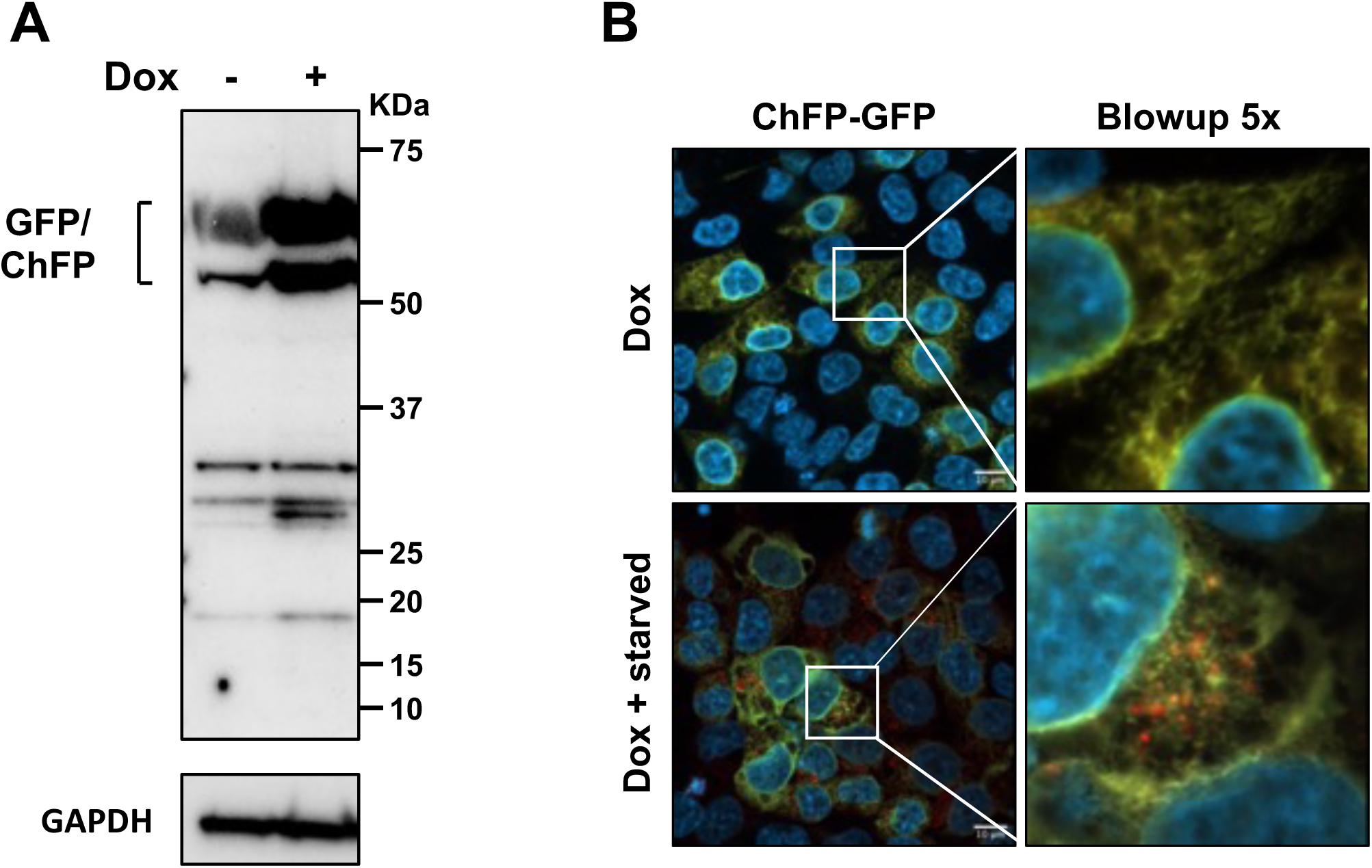
Characterization of the HCT116 er-phagy reporter cell line. A. HCT116 cells were stably transfected with and ER-phagy expressing in frame the ER-membrane resident protein RAMP4, GFP and ChFP under control of a tetracycline regulated promoter. Western blots probed with an anti-GFP antibody confirming the expression of the fusion protein in cells treated 1.5 μg/ml Dox for 24 h. B. Visualization of ER-phagy. Stably transfected HCT116 growing on cover slides were treated with Dox for 24 h followed by starvation overnight in EBSS medium. The accumulation of red dots corresponding to reporter-loaded phagolysosomes was visualized by confocal microscopy.

**Figure S5.**
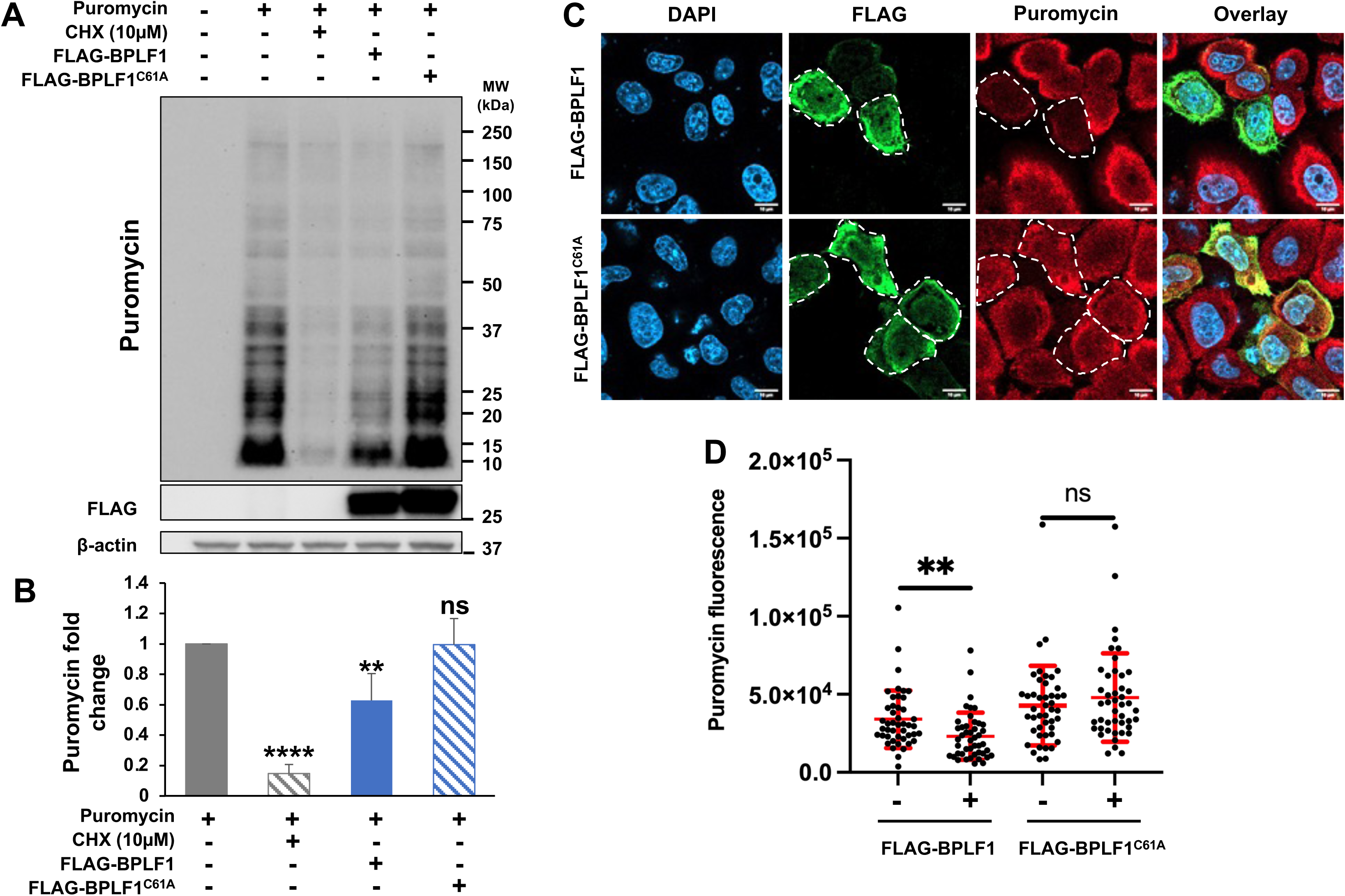
BPLF1 promotes a global inhibition of host protein synthesis. (**A**) control and FLAG-BPLF1/BPLF1^C61A^ transfected HeLa cells were cultured for 24h before analysis of protein synthesis. SUnSET assays were performed by addition of 10 µg/ml Puromycin for 10 min. As control for protein synthesis inhibition 10 μM cycloheximide (CHX) were added 5 min before Puromycin treatment. Western blots were probed with a Puromycin specific antibody. Blots from one representative experiment out four are shown in the figure. (**B**) Densitometry quantification of puromycin incorporation. The intensities of puromycin labeled bands in CHX treated and BPLF1/BPLF1^C61A^ transfected cells relative to the untreated control are shown. Mean ± SD of four independent experiments. Statistical analysis was performed using Student t-test. **P ≤ 0.01, ****P ≤ 0.0001. (**C**) SUnSET assay was performed in control and FLAG-BPLF1/BPLF1^C61A^ transfected cells grown on cover slides followed by double staining with FLAG and puromycin specific antibodies and visualization by confocal microscopy. Representative micrographs illustrating the virtual absence of puromycin fluorescence in BPLF1 cells expressing cells. Scale bar, 10 µm. (**D**) Quantification of puromycin fluorescence in BPLF1/BPLF1^C61A^ positive and negative cells from the same transfection experiment. Mean ± SD of 45 individual cells were scored for each condition. Statistical analysis was performed using Student t-test. **P ≤ 0.01, ****P ≤ 0.0001.

## Notes

### Competing Interest Statement

The authors have declared no competing interest.

## REFERENCES

Abbott, R.J., Quinn, L.L., Leese, A.M., Scholes, H.M., Pachnio, A., and Rickinson, A.B. (2013). CD8+ T cell responses to lytic EBV infection: late antigen specificities as subdominant components of the total response. J Immunol 191, 5398–5409. 10.4049/jimmunol.1301629

Ahsan, N., Kanda, T., Nagashima, K., and Takada, K. (2005). Epstein-Barr virus transforming protein LMP1 plays a critical role in virus production. J Virol 79, 4415–4424. 10.1128/JVI.79.7.4415-4424.2005

Akimitsu, N., Tanaka, J., and Pelletier, J. (2007). Translation of nonSTOP mRNA is repressed post-initiation in mammalian cells. EMBO J 26, 2327–2338. 10.1038/sj.emboj.7601679

Akopian, D., Shen, K., Zhang, X., and Shan, S.O. (2013). Signal recognition particle: an essential protein-targeting machine. Annu Rev Biochem 82, 693–721. 10.1146/annurev-biochem-072711-164732

Andreev, D.E., O’Connor, P.B., Fahey, C., Kenny, E.M., Terenin, I.M., Dmitriev, S.E., Cormican, P., Morris, D.W., Shatsky, I.N., and Baranov, P.V. (2015). Translation of 5’ leaders is pervasive in genes resistant to eIF2 repression. Elife 4, e03971. 10.7554/eLife.03971

Ashburner, M., Ball, C.A., Blake, J.A., Botstein, D., Butler, H., Cherry, J.M., Davis, A.P., Dolinski, K., Dwight, S.S., Eppig, J.T., et al. (2000). Gene ontology: tool for the unification of biology. The Gene Ontology Consortium. Nat Genet 25, 25–29. 10.1038/75556

Axten, J.M., Medina, J.R., Feng, Y., Shu, A., Romeril, S.P., Grant, S.W., Li, W.H., Heerding, D.A., Minthorn, E., Mencken, T., et al. (2012). Discovery of 7-methyl-5-(1- {[3-(trifluoromethyl)phenyl]acetyl}-2,3-dihydro-1H-indol-5-yl)-7H-p yrrolo[2,3- d]pyrimidin-4-amine (GSK2606414), a potent and selective first-in-class inhibitor of protein kinase R (PKR)-like endoplasmic reticulum kinase (PERK). J Med Chem 55, 7193–7207. 10.1021/jm300713s

Banerjee, S., Kumar, M., and Wiener, R. (2020). Decrypting UFMylation: How Proteins Are Modified with UFM1. Biomolecules 10. 10.3390/biom10101442

Barbosa, C., Peixeiro, I., and Romao, L. (2013). Gene expression regulation by upstream open reading frames and human disease. PLoS Genet 9, e1003529. 10.1371/journal.pgen.1003529

Bencun, M., Klinke, O., Hotz-Wagenblatt, A., Klaus, S., Tsai, M.H., Poirey, R., and Delecluse, H.J. (2018). Translational profiling of B cells infected with the Epstein-Barr virus reveals 5’ leader ribosome recruitment through upstream open reading frames. Nucleic Acids Res 46, 2802–2819. 10.1093/nar/gky129

Bengtson, M.H., and Joazeiro, C.A. (2010). Role of a ribosome-associated E3 ubiquitin ligase in protein quality control. Nature 467, 470–473. 10.1038/nature09371

Biswas, B., Kandpal, M., Jauhari, U.K., and Vivekanandan, P. (2016). Genome-wide analysis of G-quadruplexes in herpesvirus genomes. BMC Genomics 17, 949. 10.1186/s12864-016-3282-1

Bowman, E.J., Siebers, A., and Altendorf, K. (1988). Bafilomycins: a class of inhibitors of membrane ATPases from microorganisms, animal cells, and plant cells. Proc Natl Acad Sci U S A 85, 7972–7976. 10.1073/pnas.85.21.7972

Brandman, O., Stewart-Ornstein, J., Wong, D., Larson, A., Williams, C.C., Li, G.W., Zhou, S., King, D., Shen, P.S., Weibezahn, J., et al. (2012). A ribosome-bound quality control complex triggers degradation of nascent peptides and signals translation stress. Cell 151, 1042–1054. 10.1016/j.cell.2012.10.044

Brazeau, J.F., and Rosse, G. (2014). Triazolo[4,5-d]pyrimidine Derivatives as Inhibitors of GCN2. ACS Med Chem Lett 5, 282–283. 10.1021/ml500052f

Brink, A.A., Meijer, C.J., Nicholls, J.M., Middeldorp, J.M., and van den Brule, A.J. (2001). Activity of the EBNA1 promoter associated with lytic replication (Fp) in Epstein-Barr virus associated disorders. Mol Pathol 54, 98–102. 10.1136/mp.54.2.98

Buchan, J.R., and Stansfield, I. (2007). Halting a cellular production line: responses to ribosomal pausing during translation. Biol Cell 99, 475–487. 10.1042/BC20070037

Cai, Y., Pi, W., Sivaprakasam, S., Zhu, X., Zhang, M., Chen, J., Makala, L., Lu, C., Wu, J., Teng, Y., et al. (2015). UFBP1, a Key Component of the Ufm1 Conjugation System, Is Essential for Ufmylation-Mediated Regulation of Erythroid Development. PLoS Genet 11, e1005643. 10.1371/journal.pgen.1005643

Cesaro, T., and Michiels, T. (2021). Inhibition of PKR by Viruses. Front Microbiol 12, 757238. 10.3389/fmicb.2021.757238

Chyzynska, K., Labun, K., Jones, C., Grellscheid, S.N., and Valen, E. (2021). Deep conservation of ribosome stall sites across RNA processing genes. NAR Genom Bioinform 3, lqab038. 10.1093/nargab/lqab038

Dabral, P., Babu, J., Zareie, A., and Verma, S.C. (2020). LANA and hnRNP A1 Regulate the Translation of LANA mRNA through G-Quadruplexes. J Virol 94. 10.1128/JVI.01508-19

Daly, R., Khaperskyy, D.A., and Gaglia, M.M. (2020). Fine-tuning a blunt tool: Regulation of viral host shutoff RNases. PLoS Pathog 16, e1008385. 10.1371/journal.ppat.1008385

DiGiuseppe, S., Rollins, M.G., Bartom, E.T., and Walsh, D. (2018). ZNF598 Plays Distinct Roles in Interferon-Stimulated Gene Expression and Poxvirus Protein Synthesis. Cell Rep 23, 1249–1258. 10.1016/j.celrep.2018.03.132

Dolan, B.P., Bennink, J.R., and Yewdell, J.W. (2011). Translating DRiPs: progress in understanding viral and cellular sources of MHC class I peptide ligands. Cell Mol Life Sci 68, 1481–1489. 10.1007/s00018-011-0656-z

Donnelly, N., Gorman, A.M., Gupta, S., and Samali, A. (2013). The eIF2alpha kinases: their structures and functions. Cell Mol Life Sci 70, 3493–3511. 10.1007/s00018-012-1252-6

Ferro-Novick, S., Reggiori, F., and Brodsky, J.L. (2021). ER-Phagy, ER Homeostasis, and ER Quality Control: Implications for Disease. Trends Biochem Sci 46, 630–639. 10.1016/j.tibs.2020.12.013

Filbeck, S., Cerullo, F., Pfeffer, S., and Joazeiro, C.A.P. (2022). Ribosome-associated quality-control mechanisms from bacteria to humans. Mol Cell 82, 1451–1466. 10.1016/j.molcel.2022.03.038

Frappier, L. (2015). Ebna1. Curr Top Microbiol Immunol 391, 3–34. 10.1007/978-3-319-22834-1_1

Gak, I.A., Vasiljevic, D., Zerjatked, T., Yu, L., Brosch, M., Mourmeliotis, T.I., Horenburg, C., Klemann, N., Mabos, G., Herrmann, A., et al. (2020). UFMylation regulates translational homeostasis and cell cycle progression. bioRxív https://doi.org/10.1101/2020.02.03.931196.

Garshott, D.M., Sundaramoorthy, E., Leonard, M., and Bennett, E.J. (2020). Distinct regulatory ribosomal ubiquitylation events are reversible and hierarchically organized. Elife 9. 10.7554/eLife.54023

Garzia, A., Jafarnejad, S.M., Meyer, C., Chapat, C., Gogakos, T., Morozov, P., Amiri, M., Shapiro, M., Molina, H., Tuschl, T., et al. (2017). The E3 ubiquitin ligase and RNA-binding protein ZNF598 orchestrates ribosome quality control of premature polyadenylated mRNAs. Nat Commun 8, 16056. 10.1038/ncomms16056

Garzia, A., Meyer, C., and Tuschl, T. (2021). The E3 ubiquitin ligase RNF10 modifies 40S ribosomal subunits of ribosomes compromised in translation. Cell Rep 36, 109468. 10.1016/j.celrep.2021.109468

Gastaldello, S., Chen, X., Callegari, S., and Masucci, M.G. (2013). Caspase-1 promotes Epstein-Barr virus replication by targeting the large tegument protein deneddylase to the nucleus of productively infected cells. PLoS Pathog 9, e1003664. 10.1371/journal.ppat.1003664

Gastaldello, S., Hildebrand, S., Faridani, O., Callegari, S., Palmkvist, M., Di Guglielmo, C., and Masucci, M.G. (2010). A deneddylase encoded by Epstein-Barr virus promotes viral DNA replication by regulating the activity of cullin-RING ligases. Nat Cell Biol 12, 351–361. 10.1038/ncb2035

Gebauer, F., and Hentze, M.W. (2004). Molecular mechanisms of translational control. Nat Rev Mol Cell Biol 5, 827–835. 10.1038/nrm1488

Gerakis, Y., Quintero, M., Li, H., and Hetz, C. (2019). The UFMylation System in Proteostasis and Beyond. Trends Cell Biol 29, 974–986. 10.1016/j.tcb.2019.09.005

Gingras, A.C., Raught, B., and Sonenberg, N. (1999). eIF4 initiation factors: effectors of mRNA recruitment to ribosomes and regulators of translation. Annu Rev Biochem 68, 913–963. 10.1146/annurev.biochem.68.1.913

Gupta, S., Yla-Anttila, P., Callegari, S., Tsai, M.H., Delecluse, H.J., and Masucci, M.G. (2018). Herpesvirus deconjugases inhibit the IFN response by promoting TRIM25 autoubiquitination and functional inactivation of the RIG-I signalosome. PLoS Pathog 14, e1006852. 10.1371/journal.ppat.1006852

Houen, G., and Trier, N.H. (2020). Epstein-Barr Virus and Systemic Autoimmune Diseases. Front Immunol 11, 587380. 10.3389/fimmu.2020.587380

Huang da, W., Sherman, B.T., and Lempicki, R.A. (2009). Bioinformatics enrichment tools: paths toward the comprehensive functional analysis of large gene lists. Nucleic Acids Res 37, 1–13. 10.1093/nar/gkn923

Ikeuchi, K., Tesina, P., Matsuo, Y., Sugiyama, T., Cheng, J., Saeki, Y., Tanaka, K., Becker, T., Beckmann, R., and Inada, T. (2019). Collided ribosomes form a unique structural interface to induce Hel2-driven quality control pathways. EMBO J 38. 10.15252/embj.2018100276

Inagaki, T., Sato, Y., Ito, J., Takaki, M., Okuno, Y., Yaguchi, M., Masud, H., Watanabe, T., Sato, K., Iwami, S., et al. (2020). Direct Evidence of Abortive Lytic Infection-Mediated Establishment of Epstein-Barr Virus Latency During B-Cell Infection. Front Microbiol 11, 575255. 10.3389/fmicb.2020.575255

Isaksson, A., Berggren, M., and Ricksten, A. (2003). Epstein-Barr virus U leader exon contains an internal ribosome entry site. Oncogene 22, 572–581. 10.1038/sj.onc.1206149

Jammi, N.V., Whitby, L.R., and Beal, P.A. (2003). Small molecule inhibitors of the RNA-dependent protein kinase. Biochem Biophys Res Commun 308, 50–57. 10.1016/s0006-291x(03)01318-4

Jung, Y., Kim, H.D., Yang, H.W., Kim, H.J., Jang, C.Y., and Kim, J. (2017). Modulating cellular balance of Rps3 mono-ubiquitination by both Hel2 E3 ligase and Ubp3 deubiquitinase regulates protein quality control. Exp Mol Med 49, e390. 10.1038/emm.2017.128

Juszkiewicz, S., Chandrasekaran, V., Lin, Z., Kraatz, S., Ramakrishnan, V., and Hegde, R.S. (2018). ZNF598 Is a Quality Control Sensor of Collided Ribosomes. Mol Cell 72, 469–481 e467. 10.1016/j.molcel.2018.08.037

Juszkiewicz, S., and Hegde, R.S. (2017). Initiation of Quality Control during Poly(A) Translation Requires Site-Specific Ribosome Ubiquitination. Mol Cell 65, 743–750 e744. 10.1016/j.molcel.2016.11.039

Juszkiewicz, S., Speldewinde, S.H., Wan, L., Svejstrup, J.Q., and Hegde, R.S. (2020). The ASC-1 Complex Disassembles Collided Ribosomes. Mol Cell 79, 603–614 e608. 10.1016/j.molcel.2020.06.006

Kaimal, V., Bardes, E.E., Tabar, S.C., Jegga, A.G., and Aronow, B.J. (2010). ToppCluster: a multiple gene list feature analyzer for comparative enrichment clustering and network-based dissection of biological systems. Nucleic Acids Res 38, W96–102. 10.1093/nar/gkq418

Kelleher, D.J., and Gilmore, R. (2006). An evolving view of the eukaryotic oligosaccharyltransferase. Glycobiology 16, 47R–62R. 10.1093/glycob/cwj066

Kenney, S. (2007). Reactivation and lytic replication of EBV. In Human Herpesviruses: Biology, Therapy and Immunotherapy, A. Arvin, G. Camplanelli-Fiume, E. Mokarsky, P. Moore, B. Roizman, R. Whitley, and K. Yamanishi, eds. (Cambridge: Cambridge University press).

Kim, W., Youn, H., Lee, S., Kim, E., Kim, D., Sub Lee, J., Lee, J.M., and Youn, B. (2018). RNF138-mediated ubiquitination of rpS3 is required for resistance of glioblastoma cells to radiation-induced apoptosis. Exp Mol Med 50, e434. 10.1038/emm.2017.247

Komar, A.A., and Hatzoglou, M. (2011). Cellular IRES-mediated translation: the war of ITAFs in pathophysiological states. Cell Cycle 10, 229–240. 10.4161/cc.10.2.14472

Kwun, H.J., da Silva, S.R., Qin, H., Ferris, R.L., Tan, R., Chang, Y., and Moore, P.S. (2011). The central repeat domain 1 of Kaposi’s sarcoma-associated herpesvirus (KSHV) latency associated-nuclear antigen 1 (LANA1) prevents cis MHC class I peptide presentation. Virology 412, 357–365. 10.1016/j.virol.2011.01.026

Li, J., Nagy, N., Liu, J., Gupta, S., Frisan, T., Hennig, T., Cameron, D.P., Baranello, L., and Masucci, M.G. (2021). The Epstein-Barr virus deubiquitinating enzyme BPLF1 regulates the activity of topoisomerase II during productive infection. PLoS Pathog 17, e1009954. 10.1371/journal.ppat.1009954

Liang, J.R., Lingeman, E., Ahmed, S., and Corn, J.E. (2018). Atlastins remodel the endoplasmic reticulum for selective autophagy. J Cell Biol 217, 3354–3367. 10.1083/jcb.201804185

Liang, J.R., Lingeman, E., Luong, T., Ahmed, S., Muhar, M., Nguyen, T., Olzmann, J.A., and Corn, J.E. (2020). A Genome-wide ER-phagy Screen Highlights Key Roles of Mitochondrial Metabolism and ER-Resident UFMylation. Cell 180, 1160–1177 e1120. 10.1016/j.cell.2020.02.017

Liu, J., Guan, D., Dong, M., Yang, J., Wei, H., Liang, Q., Song, L., Xu, L., Bai, J., Liu, C., et al. (2020). UFMylation maintains tumour suppressor p53 stability by antagonizing its ubiquitination. Nat Cell Biol 22, 1056–1063. 10.1038/s41556-020-0559-z

Liu, J., Wang, Y., Song, L., Zeng, L., Yi, W., Liu, T., Chen, H., Wang, M., Ju, Z., and Cong, Y.S. (2017). A critical role of DDRGK1 in endoplasmic reticulum homoeostasis via regulation of IRE1alpha stability. Nat Commun 8, 14186. 10.1038/ncomms14186

Lu, P.D., Harding, H.P., and Ron, D. (2004). Translation reinitiation at alternative open reading frames regulates gene expression in an integrated stress response. J Cell Biol 167, 27–33. 10.1083/jcb.200408003

Masucci, M.G. (2022). Herpesvirus ubiquitin deconjugases. Semin Cell Dev Biol 132, 185–192. 10.1016/j.semcdb.2021.10.011

Masutani, M., Sonenberg, N., Yokoyama, S., and Imataka, H. (2007). Reconstitution reveals the functional core of mammalian eIF3. EMBO J 26, 3373–3383. 10.1038/sj.emboj.7601765

Matsuo, Y., Ikeuchi, K., Saeki, Y., Iwasaki, S., Schmidt, C., Udagawa, T., Sato, F., Tsuchiya, H., Becker, T., Tanaka, K., et al. (2017). Ubiquitination of stalled ribosome triggers ribosome-associated quality control. Nat Commun 8, 159. 10.1038/s41467-017- 00188-1

Matsuo, Y., Tesina, P., Nakajima, S., Mizuno, M., Endo, A., Buschauer, R., Cheng, J., Shounai, O., Ikeuchi, K., Saeki, Y., et al. (2020). RQT complex dissociates ribosomes collided on endogenous RQC substrate SDD1. Nat Struct Mol Biol 27, 323–332. 10.1038/s41594-020-0393-9

Meyer, C., Garzia, A., Morozov, P., Molina, H., and Tuschl, T. (2020). The G3BP1- Family-USP10 Deubiquitinase Complex Rescues Ubiquitinated 40S Subunits of Ribosomes Stalled in Translation from Lysosomal Degradation. Mol Cell 77, 1193–1205 e1195. 10.1016/j.molcel.2019.12.024

Munz, C. (2021). The Role of Lytic Infection for Lymphomagenesis of Human gamma- Herpesviruses. Front Cell Infect Microbiol 11, 605258. 10.3389/fcimb.2021.605258

Murat, P., Zhong, J., Lekieffre, L., Cowieson, N.P., Clancy, J.L., Preiss, T., Balasubramanian, S., Khanna, R., and Tellam, J. (2014). G-quadruplexes regulate Epstein-Barr virus-encoded nuclear antigen 1 mRNA translation. Nat Chem Biol 10, 358–364. 10.1038/nchembio.1479

Nunes, A., Ribeiro, D.R., Marques, M., Santos, M.A.S., Ribeiro, D., and Soares, A.R. (2020). Emerging Roles of tRNAs in RNA Virus Infections. Trends Biochem Sci 45, 794–805. 10.1016/j.tibs.2020.05.007

Osborne, A.R., Rapoport, T.A., and van den Berg, B. (2005). Protein translocation by the Sec61/SecY channel. Annu Rev Cell Dev Biol 21, 529–550. 10.1146/annurev.cellbio.21.012704.133214

Pelechano, V., and Alepuz, P. (2017). eIF5A facilitates translation termination globally and promotes the elongation of many non polyproline-specific tripeptide sequences. Nucleic Acids Res 45, 7326–7338. 10.1093/nar/gkx479

Peter, J., Magnussen, H., DeRosa, P., Millrine, D., Mattews, S., Lamoliatte, F., Sundaramoorthy, R., Kopito, R., and Kulathu, Y. (2022). Non canonical Scaffols-type ligase complex mediates protein UFMylation. bioRxív https://doi.org/10.1101/2022.01.31.478489.

Qin, B., Yu, J., Nowsheen, S., Wang, M., Tu, X., Liu, T., Li, H., Wang, L., and Lou, Z. (2019). UFL1 promotes histone H4 ufmylation and ATM activation. Nat Commun 10, 1242. 10.1038/s41467-019-09175-0

Ruggiano, A., Foresti, O., and Carvalho, P. (2014). Quality control: ER-associated degradation: protein quality control and beyond. J Cell Biol 204, 869–879. 10.1083/jcb.201312042

Sehgal, P., Szalai, P., Olesen, C., Praetorius, H.A., Nissen, P., Christensen, S.B., Engedal, N., and Moller, J.V. (2017). Inhibition of the sarco/endoplasmic reticulum (ER) Ca(2+)-ATPase by thapsigargin analogs induces cell death via ER Ca(2+) depletion and the unfolded protein response. J Biol Chem 292, 19656–19673. 10.1074/jbc.M117.796920

Shannon, P., Markiel, A., Ozier, O., Baliga, N.S., Wang, J.T., Ramage, D., Amin, N., Schwikowski, B., and Ideker, T. (2003). Cytoscape: a software environment for integrated models of biomolecular interaction networks. Genome Res 13, 2498–2504. 10.1101/gr.1239303

Shannon-Lowe, C., and Rickinson, A. (2019). The Global Landscape of EBV-Associated Tumors. Front Oncol 9, 713. 10.3389/fonc.2019.00713

Shao, S., von der Malsburg, K., and Hegde, R.S. (2013). Listerin-dependent nascent protein ubiquitination relies on ribosome subunit dissociation. Mol Cell 50, 637–648. 10.1016/j.molcel.2013.04.015

Sidrauski, C., Acosta-Alvear, D., Khoutorsky, A., Vedantham, P., Hearn, B.R., Li, H., Gamache, K., Gallagher, C.M., Ang, K.K., Wilson, C., et al. (2013). Pharmacological brake-release of mRNA translation enhances cognitive memory. Elife 2, e00498. 10.7554/eLife.00498

Sivachandran, N., Wang, X., and Frappier, L. (2012). Functions of the Epstein-Barr virus EBNA1 protein in viral reactivation and lytic infection. J Virol 86, 6146–6158. 10.1128/JVI.00013-12

Snel, B., Lehmann, G., Bork, P., and Huynen, M.A. (2000). STRING: a web-server to retrieve and display the repeatedly occurring neighbourhood of a gene. Nucleic Acids Res 28, 3442–3444. 10.1093/nar/28.18.3442

Song, J., Perreault, J.P., Topisirovic, I., and Richard, S. (2016). RNA G-quadruplexes and their potential regulatory roles in translation. Translation (Austin) 4, e1244031. 10.1080/21690731.2016.1244031

Starck, S.R., Tsai, J.C., Chen, K., Shodiya, M., Wang, L., Yahiro, K., Martins-Green, M., Shastri, N., and Walter, P. (2016). Translation from the 5’ untranslated region shapes the integrated stress response. Science 351, aad3867. 10.1126/science.aad3867

Stephani, M., Picchianti, L., and Dagdas, Y. (2021). C53 is a cross-kingdom conserved reticulophagy receptor that bridges the gap betweenselective autophagy and ribosome stalling at the endoplasmic reticulum. Autophagy 17, 586–587. 10.1080/15548627.2020.1846304

Sundaramoorthy, E., Leonard, M., Mak, R., Liao, J., Fulzele, A., and Bennett, E.J. (2017). ZNF598 and RACK1 Regulate Mammalian Ribosome-Associated Quality Control Function by Mediating Regulatory 40S Ribosomal Ubiquitylation. Mol Cell 65, 751–760 e754. 10.1016/j.molcel.2016.12.026

Sundaramoorthy, E., Ryan, A.P., Fulzele, A., Leonard, M., Daugherty, M.D., and Bennett, E.J. (2021). Ribosome quality control activity potentiates vaccinia virus protein synthesis during infection. J Cell Sci 134. 10.1242/jcs.257188

Tatsumi, K., Sou, Y.S., Tada, N., Nakamura, E., Iemura, S., Natsume, T., Kang, S.H., Chung, C.H., Kasahara, M., Kominami, E., et al. (2010). A novel type of E3 ligase for the Ufm1 conjugation system. J Biol Chem 285, 5417–5427. 10.1074/jbc.M109.036814

The Gene Ontology, C. (2019). The Gene Ontology Resource: 20 years and still GOing strong. Nucleic Acids Res 47, D330–D338. 10.1093/nar/gky1055

Verma, R., Oania, R.S., Kolawa, N.J., and Deshaies, R.J. (2013). Cdc48/p97 promotes degradation of aberrant nascent polypeptides bound to the ribosome. Elife 2, e00308. 10.7554/eLife.00308

Vind, A.C., Genzor, A.V., and Bekker-Jensen, S. (2020). Ribosomal stress-surveillance: three pathways is a magic number. Nucleic Acids Res 48, 10648–10661. 10.1093/nar/gkaa757

Walczak, C.P., Leto, D.E., Zhang, L., Riepe, C., Muller, R.Y., DaRosa, P.A., Ingolia, N.T., Elias, J.E., and Kopito, R.R. (2019). Ribosomal protein RPL26 is the principal target of UFMylation. Proc Natl Acad Sci U S A 116, 1299–1308. 10.1073/pnas.1816202116

Wang, L., Xu, Y., Rogers, H., Saidi, L., Noguchi, C.T., Li, H., Yewdell, J.W., Guydosh, N.R., and Ye, Y. (2020). UFMylation of RPL26 links translocation-associated quality control to endoplasmic reticulum protein homeostasis. Cell Res 30, 5–20. 10.1038/s41422-019-0236-6

Wilkinson, S. (2019). ER-phagy: shaping up and destressing the endoplasmic reticulum. FEBS J 286, 2645–2663. 10.1111/febs.14932

Witting, K.F., and Mulder, M.P.C. (2021). Highly Specialized Ubiquitin-Like Modifications: Shedding Light into the UFM1 Enigma. Biomolecules 11. 10.3390/biom11020255

Wu, C.C., Peterson, A., Zinshteyn, B., Regot, S., and Green, R. (2020). Ribosome Collisions Trigger General Stress Responses to Regulate Cell Fate. Cell 182, 404–416 e414. 10.1016/j.cell.2020.06.006

Yan, L.L., and Zaher, H.S. (2021). Ribosome quality control antagonizes the activation of the integrated stress response on colliding ribosomes. Mol Cell 81, 614–628 e614. 10.1016/j.molcel.2020.11.033

Zhang, M., Zhu, X., Zhang, Y., Cai, Y., Chen, J., Sivaprakasam, S., Gurav, A., Pi, W., Makala, L., Wu, J., et al. (2015). RCAD/Ufl1, a Ufm1 E3 ligase, is essential for hematopoietic stem cell function and murine hematopoiesis. Cell Death Differ 22, 1922–1934. 10.1038/cdd.2015.51

## References

van Gent, M., Braem, S.G., de Jong, A., Delagic, N., Peeters, J.G., Boer, I.G., Moynagh, P.N., Kremmer, E., Wiertz, E.J., Ovaa, H., et al. (2014). Epstein-Barr virus large tegument protein BPLF1 contributes to innate immune evasion through interference with toll-like receptor signaling. PLoS Pathog 10, e1003960. 10.1371/journal.ppat.1003960

